# A reservoir of rituximab-resistant splenic memory B cells contributes to relapses after B-cell depletion therapy

**DOI:** 10.1101/833343

**Authors:** Etienne Crickx, Pascal Chappert, Sandra Weller, Aurélien Sokal, Imane Azzaoui, Alexis Vandenberghe, Guillaume Bonnard, Geoffrey Rossi, Tatiana Fadeev, Sébastien Storck, Lionel Galicier, Véronique Meignin, Etienne Rivière, Bertrand Godeau, Marc Michel, Jean-Claude Weill, Claude-Agnès Reynaud, Matthieu Mahévas

**Author notes:** Address correspondence: Dr Matthieu Mahévas, Service de médecine interne, Hôpital Henri-Mondor, Assistance Publique-Hôpitaux de Paris (AP-HP), Créteil.

## Abstract

Immune thrombocytopenia (ITP) is an autoimmune disease mediated by pathogenic antibodies directed against platelet antigens, including GPIIbIIIa. Taking advantage of spleen samples obtained from ITP patients, we characterized by multiples approaches the onset of disease relapses occurring after an initial complete response to rituximab. Analysis of splenic B cell immunoglobulin heavy chain gene repertoire at bulk level and from single anti-GPIIbIIIa B cells revealed that germinal centers were fueled by B cells originating from the ongoing lymphopoiesis, but also by rituximab-resistant memory B cells, both giving rise to anti-GPIIbIIIa plasma cells. We identified a population of splenic memory B cells that resisted rituximab through acquisition of a unique phenotype and contributed to relapses, providing a new target in B cell mediated autoimmune diseases.

## Introduction

Primary immune thrombocytopenia (ITP) is a prototypic B cell mediated autoimmune disease in which pathogenic antibodies directed against the platelet membrane glycoprotein IIb-IIIa (GPIIbIIIa) lead to platelet destruction by macrophages in the spleen (Kuwana et al., 1998, 2009). In active ITP patients, the spleen is the site of an intense autoimmune response with germinal center (GC) expansion, resulting in the secretion of mutated anti-GPIIbIIIa antibodies by plasma cells (PC) (Mahévas et al., 2012; Roark et al., 2002). While these anti-platelet PC have been shown to play a central role in ITP, the contribution of autoreactive memory B cells to the disease pathogeny remains elusive, as for many antibody-mediated auto-immune diseases.

B cell depletion with the anti-CD20 chimeric antibody rituximab (RTX) has constituted a major step forward in the treatment of antibody mediated auto-immune diseases, but patients achieving an initial complete response frequently relapse during B-cell reconstitution. The fundamental mechanisms underlying this clinical observation are not well understood. In ITP, B-cell depletion with RTX has an immediate response rate of 50%, and patients failing to respond or relapsing are subsequently splenectomized (Provan et al., 2010). Therapeutic splenectomy, resulting in a durable platelet response in 60-70 % of ITP patients, offers a unique access to study the autoimmune response in the spleen. Despite the fact that complete B-cell depletion is achieved in peripheral blood in most patients receiving RTX, we and others have found evidence for a residual CD19+ B cell population in the spleen of primary failure patients (i.e: splenectomized 1-6 months after RTX), mainly represented by PC and memory B cells (Audia et al., 2011; Mahévas et al., 2012). Moreover, we demonstrated that B-cell depletion per se, via its impact on the splenic microenvironment, could promote the emergence of long-lived plasma cells (LLPCs) in the spleen, some of them being autoreactive, thus explaining the primary failure of RTX in some patients (Mahévas et al., 2012). In contrast, patients achieving a complete response after RTX have most probably no remaining anti-platelet LLPCs during clinical remission. Nevertheless, a large proportion of these patients eventually relapse during B cell reconstitution (Khellaf et al., 2014; Patel et al., 2012) when RTX is cleared from the organism (usually 6 months after the last infusion (Lioger et al., 2017; Ng et al., 2005)).

Two main scenarios could explain disease relapses: either a new tolerance breakdown could promote the recruitment of newly formed naïve B cells in the autoimmune response, or alternatively, residual autoreactive memory B cells that survived RTX could be reactivated. This latter hypothesis nevertheless implies that RTX-resistant memory B cells remained quiescent during months despite the persistence of platelet antigens.

Here, taking advantage of spleen samples obtained from ITP patients, we characterized by multiple approaches the onset of disease relapses occurring after an initial complete response to RTX. We showed that germinal centers giving rise to autoreactive PC are fueled by newly generated B cells as well as by RTX-resistant memory B cells. This latter population resisted RTX through acquisition of a unique phenotype and contributed to relapses, providing a new therapeutic opportunity in B cell mediated autoimmune diseases.

## Results

### Patients

We analyzed spleen samples from 8 patients (median age 45.5 years, range 30 – 65) that achieved a complete response after a course of RTX (2 doses of 1g at day 1 and 15, or 375 mg/m² at day 1, 8, 15 and 22) and relapsed 8.5 months in median (range 5 – 13) after the last infusion (hereafter referred to as “RTX relapse” patients). Clinical characteristics of patients are presented in **Supplementary Table 1**. Platelet counts were normal during remission, suggesting that the number of remaining plasma cells producing antibodies directed against platelets was not significant and that relapses were caused by the differentiation of either activated naïve B cells and/or memory B cells into autoreactive plasma cells. We also analyzed 7 patients with active ITP that were not treated with RTX (“ITP” patients), 9 healthy donors with no immune disease that died from stroke (“HD” patients) as controls and 16 patients that received RTX 1 to 6 months before splenectomy (“RTX failure” patients) and had not reconstituted their B cell pool yet.

### Spleens of RTX relapse patients reveal synchronous B cell reconstitution and GC-derived autoimmune response leading to the generation of IgG anti-GPIIbIIIa plasma cells

We investigated the presence of B cell subpopulations in spleen from RTX relapse patients (**Figure 1A**). B-cell reconstitution had started in all patients, albeit CD19+ cells were present in lower proportions compared to HD and ITP patients. CD19+ cells were mainly naïve B cells (**Figure 1C**), with an increased proportion of transitional B cells compared to HD and ITP patients (**Figure 1D**). By contrast, switched memory B-cells (**Figure 1E**) and marginal zone B-cells (**Figure 1F**) were significantly decreased compared to HD and ITP, while double negative CD27-IgD-(**Figure 1G**) appeared unchanged, suggesting that even if a new wave of transitional and naïve B cells had started to repopulate the spleen, the reconstitution of the memory pool had barely begun.

**Figure 1:**
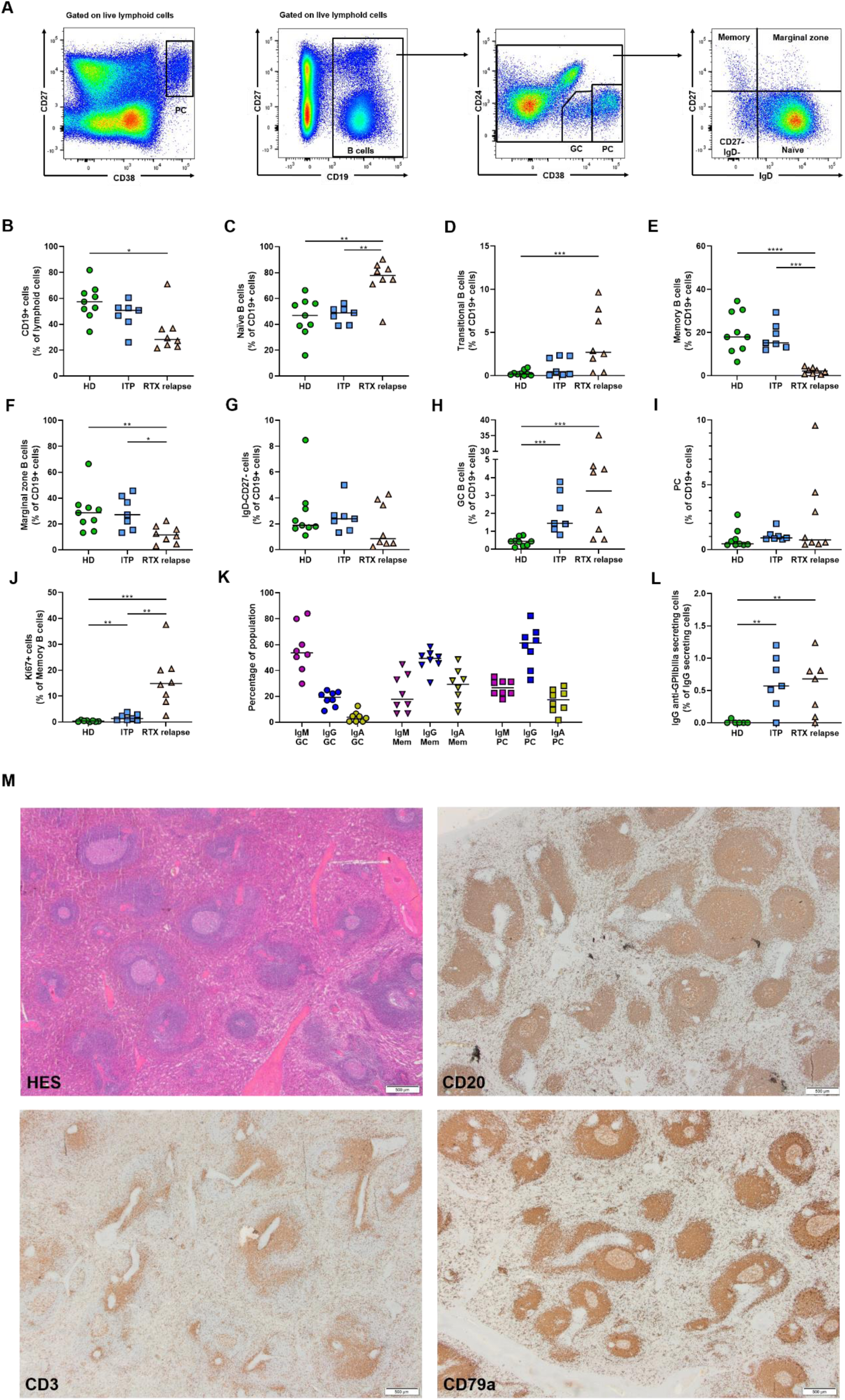
GC derived autoimmune response leading to the generation of IgG anti-GPIIbIIIa plasma cells in RTX relapse. **(A)** Representative dot plots showing gating strategy for flow cytometry analysis of splenic mononuclear cells labeled with antibodies to CD19, CD24, CD27, CD38, IgD. PC were identified as CD27^hi^CD38^hi^ cells among live lymphoid cells. After gating on CD19+ cells among live lymphoid cells, GC B cells were identified as CD24–CD38^int^. After exclusion of GC and PC, B cells were further subdivided into memory B cells (CD27+IgD-), double negative (CD27-IgD-), marginal zone B cells (CD27+IgD+), and naïve B cells (CD27-, IgD+). (**B-J**) Proportion of splenic B cells **(B)**, naïve B cells **(C)**, transitional B cells (identified as CD19+CD27-IgD+CD24hiCD38hiCD10+) **(D)**, memory B cells **(E)**, marginal zone B cells **(F)**, CD27-IgD-B cells **(G)**, GC B cells **(H)**, PC **(I)**, Ki67+ memory B cells **(J)**, in HD (n=9), ITP patients (n=7) and RTX relapse patients (n=8). (**K**) Proportion of IgM, IgG and IgA expressed by splenic GC B cells, memory B cell and PC assessed by flow cytometry analysis after intracellular staining in RTX relapse patients (n=8). (**L**) Frequency of IgG anti-GPIIbIIIa secreting cells among IgG secreting cells assessed by ELISPOT assay in HD (n=6), ITP patients (n=7) and RTX relapse patients (n=7). Two-tailed Mann-Whitney tests were performed (\*\*\**P* < 0.001; \*\**P* < 0.01, \**P* < 0.05). Symbols indicate individual samples; horizontal bars represent median values. **(M)** Representative spleen sections from RTX relapse patients (n=2) stained with hematoxylin and eosin, CD20, CD3 and CD79a. Scale bars: 500 μm.

GC B cell population was significantly and largely expanded in RTX relapse patients compared to HD, suggesting that, as for ITP patients, these structures were central in the ongoing autoreactive immune response (**Figure 1H**). We confirmed by microscopy analysis the presence of numerous CD20+ GC structures within secondary follicles, PD1 T-follicular helper cells and CD21 follicular dendritic cells (**Figure 1 and Supplementary Figure 1**). GC B cells were mainly composed of IgM+ cells, while IgG+ cells represented only a small fraction and the proportion of IgA+ cells was negligible (**Figure 1K**). Interestingly, we observed a significant proportion of Ki67 expressing memory B cells in RTX relapse patients (**Figure 1J**), but not in ITP and HD patients. Proliferating memory B cells were almost only of IgM and IgG isotypes (data not shown), arguing against a mucosal origin of these cells, and in accordance with the rarity of IgA+ cells in GC. This suggested that, despite their low numbers, memory B-cells were involved in an active auto-immune response. Next, we wondered if the germinal center response resulted in the production of autoreactive plasma cells. Steroids given before splenectomy in ITP and RTX relapse patients may have reduced the proportion of plasma cells, which were similar between all patient groups. Despite this, we observed approximately 1% of anti-GpIIbIIIa IgG-secreting cells in RTX relapse patients, a proportion similar to those observed in ITP patients (**Figure 1L**).

Overall, we found that B cell reconstitution in the spleen of RTX relapse patients was characterized by a high proportion of newly generated naïve and transitional B cells while the memory B cell pool remained severely reduced, similarly to previous findings in peripheral blood. However, we found evidences for an ongoing autoimmune response involving germinal center structures and leading to the generation of IgG anti-GPIIbIIIa plasma cells.

### Analysis of splenic B cell IgH repertoire reveals the coexistence of RTX-resistant and newly generated B cells

Clonal expansions in germinal centers give rise to clonally related memory B cells and plasma cells. We analyzed the splenic B cell repertoire of 3 RTX relapse patients and sorted PC (CD19+CD27hiCD38hi), GC (CD19+CD24-CD38intIgD-CD20+), memory (CD19+CD24+CD27+CD38-IgD-) and naïve (CD19+CD27-CD38-/intIgD+) B cells to perform high throughput sequencing of immunoglobulin heavy chain genes (**Supplementary Table 2**). As expected, VH and JH usage and CDR3 length were different between naïve and antigen-experienced B cells (**Supplementary Figure 2A, 2B and 2C**). Clonal expansions of variable sizes were found in GC, memory and PC populations (**Supplementary Figure 2D**). Because IgA expression was minor among GC B cells, we focused our analysis on IgM and IgG sequences.

Analysis of VH segment mutation distribution revealed that a large fraction of sequences had acquired few mutations (**Figure 2**), consistent with the presence of newly activated naïve B cells joining the GC. Surprisingly, another peak of highly mutated GC sequences was observed, the GC sequence pool thus harboring a bimodal distribution (**Figure 2**). Analysis of VH mutation in IgM and IgG sequences confirmed that unmutated/low mutated sequences were mostly IgM, while there was a predominance of IgG in highly mutated sequences (**Supplementary Figure 3A**). Interestingly, these bimodal distributions mirrored closely those observed in PC populations, where some cells resisted RTX while some others were generated through GC reactions. Because it is very unlikely that B cells and PCs accumulated up to 30-40 mutations during the few weeks following B cell reconstitution and relapse, this strongly suggested that low mutated sequences originated from newly generated B cells, while highly mutated sequences originated from memory B cells and PCs that escaped RTX depletion.

**Figure 2:**
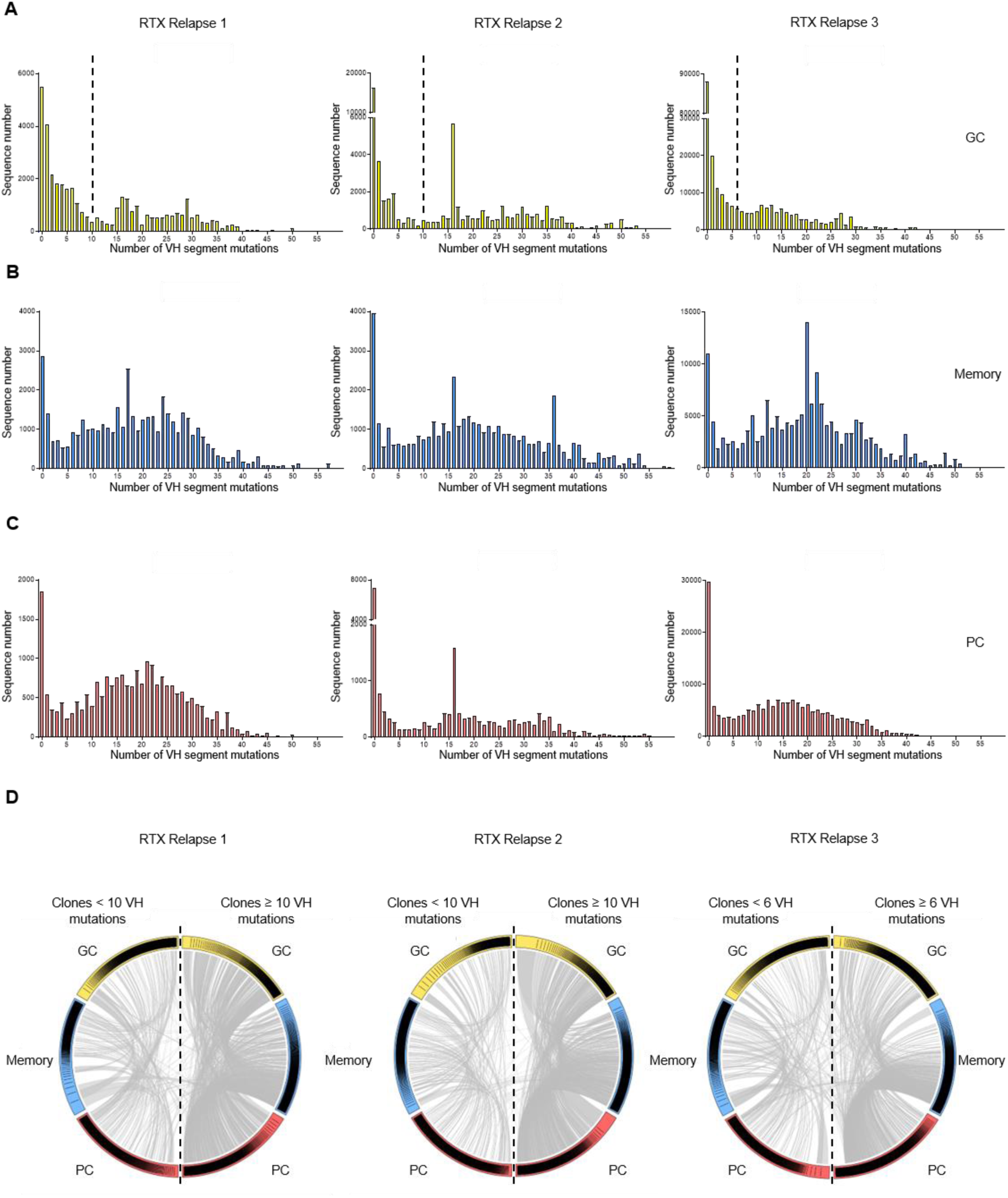
Rituximab-resistant and newly generated B cells coexist in the spleen of RTX relapse patients. **(A-C)** VH segment mutation distribution in IgM and IgG sequences from **(A)** splenic GC (yellow), **(B)** memory (blue) and **(C)** PC (red) populations from 3 RTX relapse patients was assessed by high throughput IgH sequencing. All subpopulations displayed a bimodal distribution with 2 peaks corresponding to unmutated/low mutated cells, and highly mutated cells. Dotted lines indicate the threshold used in **(D)** to discriminate low mutated clones and highly mutated clones. **(D)** Circos plot showing clonal relationships shared between IgM and IgG sequences from GC, memory and PC splenic populations. According to the mutation distribution in GC shown in **(A)**, clones from each population were classified into “low mutated” (left side of the plot) and “highly mutated” (right side of the plot) based on the clone median mutation number. Each colored sector represents one subpopulation and is divided into segments representing individual clones in rank-sized order. The internal connections show clones common to multiple subpopulations. Few clonal relationships were found between low and highly mutated populations, suggesting a distinct origin.

To confirm that low mutated and highly mutated sequences indeed originated from distinct populations, we categorized clones in either low mutated or highly mutated populations according to a cut-off value extrapolated from the mutation distribution in GC (i.e: ≥ 10 or < 10 mutations for patients 1 and 2, and ≥ 6 or < 6 for patients 3). This dichotomy was biologically relevant, because mutation distribution inside each clone showed limited dispersion (standard deviation in mutation frequency for the first 100 clones is shown in **Supplementary Table 3**). We then analyzed clonal relationships shared between GC, memory and PC clones in both populations (**Figure 2B and Supplementary Figure 3B**). Numerous clones were present in GC, memory and PC subsets in either low mutated or highly mutated populations. Among the total number of clonal relationships, only 5.7%, 3.5 and 11.5% were shared between low mutated and highly mutated populations for patients 1, 2 and 3, respectively. Moreover, most of these clones had a median mutation number that was close to the cut-off value selected for our analysis.

These results confirmed that low mutated and highly mutated clones were distinct populations, suggesting that both activated, newly generated naïve B cells and RTX-resistant memory B cells were able to fuel GC reactions and to give rise to PC in RTX relapse patients.

### GPIIbIIIa specific B cells are found in both RTX-resistant and newly generated B cells populations

Next, we wondered if autoreactive cells were enriched in RTX resistant or newly generated B cell populations. We sorted single GC and IgG+ memory B cells from the spleen of RTX relapse patients and cultured them in plates coated with CD40L expressing cells with BAFF, IL2, IL4 and IL21 using the Nojima culture system (McCarthy et al., 2018; Nojima et al., 2011). These conditions allowed for GC B cells survival and immunoglobulin class-switch *in vitro*, as confirmed by detectable IgM and/or IgG in supernatants (data not shown). GC and memory B cells proliferated and differentiated into plasmablasts *in vitro*, allowing for testing their Ag specificity. Because IgG concentrations were highly variable from one well to another, we used ELISPOT assays to identify GPIIbIIIa-specific clones and to obtain specific IgH sequences (**Figure 3A**). This method had already been validated for GPIIbIIIa identification of IgG secreting cells and we confirmed that the AP3 mouse hybridoma (secreting anti-GPIIIa antibodies) was correctly detected even at very low cell numbers (**Supplementary Figure 4A**). To confirm the anti-GPIIbIIIa specificity, we cloned IgH and IgL genes from 2 GPIIbIIIa ELISPOT-positive memory B cells and re-expressed them in HEK cells (**Supplementary Figure 4B**). Supernatants from both clones reacted with GPIIbIIIa in ELISA and Monoclonal Antibody Immobilization of Platelet Antigens (MAIPA, reference method for anti-GPIIbIIIa antibody detection in the clinic) assays.

**Figure 3:**
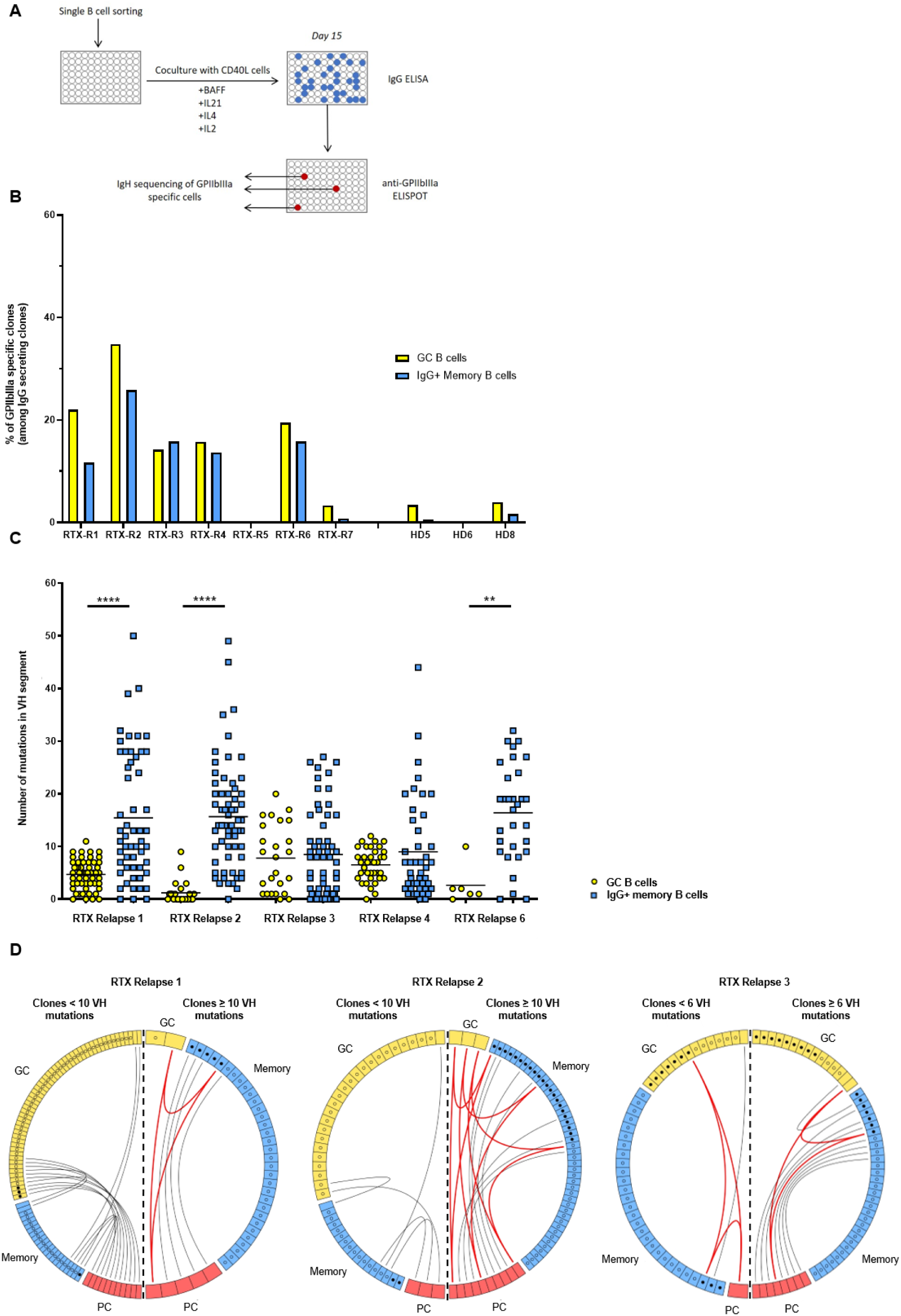
GPIIbIIIa specific B cells are found in both rituximab resistant and newly generated B cells populations. **(A)** Identification of GPIIbIIIa-specific B cells. Single GC and memory B cells were directly sorted into 96 wells plates containing CD40L expressing cells and cytokine cocktail and grown for 15 days. IgG ELISA was then performed on supernatants to select IgG secreting clones for GPIIbIIIa ELISPOT. Cells from GPIIbIIIa-specific clones were collected for IgH sequencing. **(B)** Frequency of GPIIbIIIa-specific clones among GC (yellow) and IgG memory B cells (blue) populations in RTX relapse patients and healthy donors. The total number of tested cells is given in supplementary Table 2. RTX relapse patient 8 was not tested because of low culture yield. **(C)** Number of mutations in VH segment for individual GPIIbIIIa specific GC (yellow) and IgG memory (blue) B cells from RTX relapse patients #1, #2, #3, #4 and #6. Two-tailed Mann-Whitney tests (\*\*\*\**P* < 0.0001; \*\*\**P* < 0.001; \*\**P* < 0.01, \**P* < 0.05). Symbols indicate individual cells; horizontal bars represent mean values. **(D)** Circos plot showing clonal relationships between individual sequences from GPIIbIIIa specific GC or IgG+ memory B cells and sequences obtained from high-throughput IgH analysis of RTX relapse patients #1, #2 and #3. Each colored sector represents one subpopulation and is divided into segments representing individual clones. Each circle represents a sequence originating from GPIIbIIIa B cells and is filled with black if the sequence was found in high throughput IgH sequencing data from the same subpopulation. The internal connections show clones shared by two or three subpopulations, depicted in grey or red, respectively. According to the mutation distribution in GC shown in **Figure 2**, clones from each population were classified into “low mutated” (left side of the plot) and “highly mutated” (right side of the plot) based on the clone mutation number (median clone mutation number was used for clones from high throughput IgH sequencing data). Only clonal relationships shared between individual sequences and high-throughput IgH analysis data are shown.

Using this system, 5 out of 7 RTX relapse patients had anti-GPIIbIIIa specific cells above background level. The proportion of GPIIbIIIa-specific cells ranged from 14 to 35% in GC and from 12 to 26% in IgG+ memory B cells in these patients (**Figure 3 and Supplementary Table 4**). We then analyzed the IgH sequences from these single anti-GPIIbIIIa specific cells. The repertoire of VH genes was diverse, VH3-23 and VH3-30 being the most represented ones (**Supplementary Figure 5**). GPIIbIIIa specific GC sequences had few VH segment mutations, suggesting that most of these cells originated from newly generated B cells. A low mutation count was also found in some memory B cell sequences, probably originating from newly generated B cells in GC. In contrast, all patients had IgG+ memory B cell sequences containing a high number of mutations, suggesting that most of them had anti-GPIIbIIIa RTX resistant memory B cells.

To confirm that GPIIbIIIa specific cells identified with our system were indeed implicated in an active immune response, we searched for shared clones in data from IgH high throughput sequencing (**Figure 3**). We found that some highly mutated sequences from GPIIbIIIa-specific IgG+ memory B cells belonged to clones found in GC and/or PC populations, suggesting that autoreactive, RTX-resistant memory B cells were able to give rise to autoreactive PC either directly or after returning into GC (**Figure 3 and Figure 4**). We also found share GPIIbIIIa-specific clones with few VH mutations in GC and/or memory and/or PC populations, suggesting that an ongoing immune response involving newly generated B cells also coexisted in all patients.

**Figure 4:**
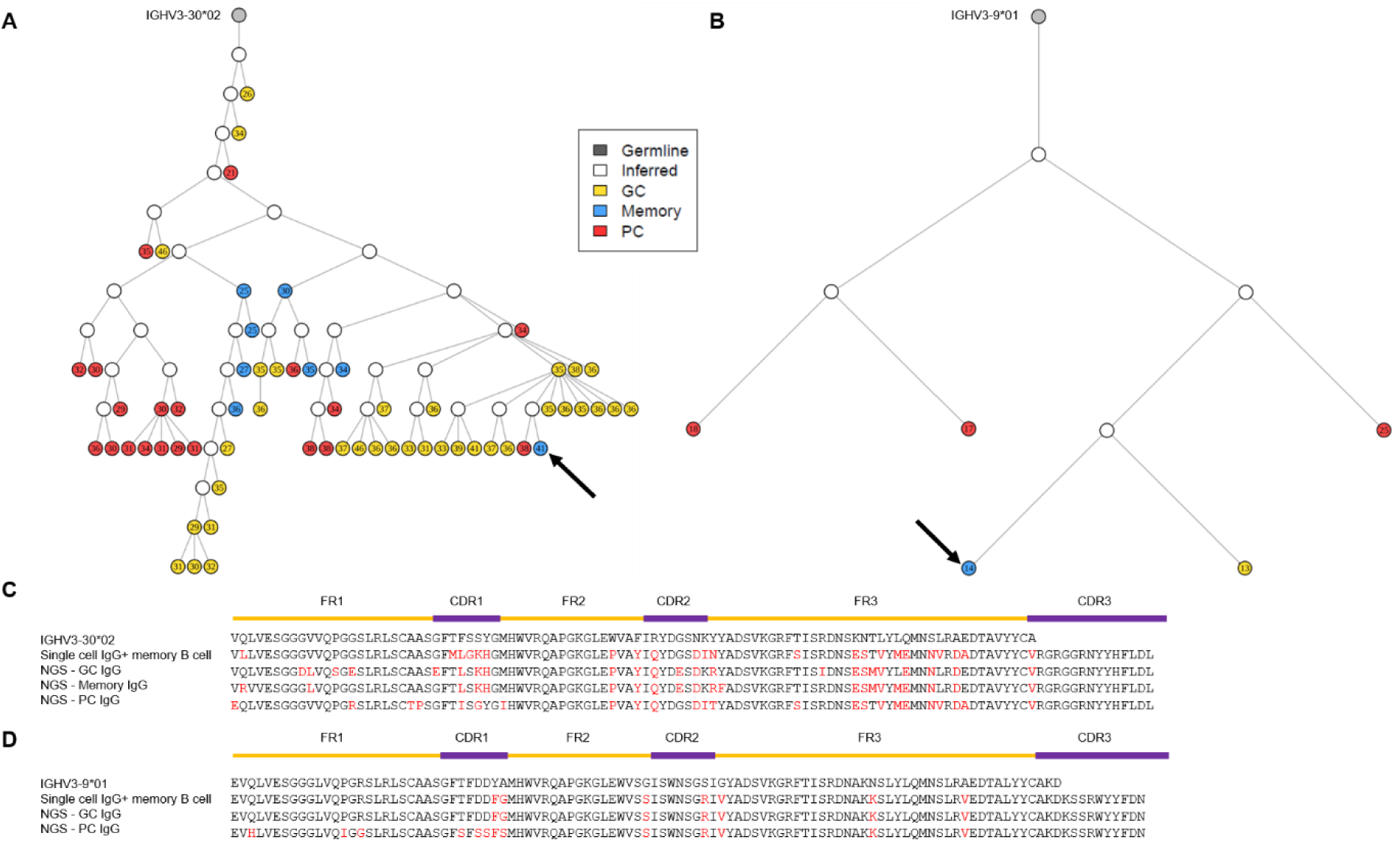
Highly mutated anti-GPIIbIIIa clones are found in GC, MS and PC populations. Phylogenetic analysis of representative clones from the high throughput IgH sequencing data containing anti-GPIIbIIIa B cells from patient 1 **(A)** and 2 **(B)**. Germline sequences are represented in grey, GC in yellow, memory B cells in blue, PC in red and open circles represent inferred precursors. The number of mutations in VH segment is indicated in each circle. All sequences are of IgG isotype. Arrows indicate specific anti-GPIIbIIIa sequences from single cell assay. **(C)** and **(D)**, representative sequence alignment of the same clones; amino-acid changes from germline are shown in red.

Overall, these findings revealed that splenic GC from RTX relapse patients are enriched in anti-GPIIbIIIa cells and are composed mainly of newly generated B cells, but also of highly mutated cells originating from RTX-resistant memory B cells. Interestingly, both populations were able to differentiate into anti-GPIIbIIIa PC, thus contributing to disease relapses.

### RTX-resistant memory B cells have unique surface phenotype and contain autoreactive clones

To confirm the existence of RTX resistant memory B cells and characterize their phenotype and function, we analyzed spleen samples from 16 patients that received RTX 1 to 6 months before splenectomy (RTX failure patients) and had not reconstituted their B cell pool yet (**Supplementary Table 1**). In these spleens, CD19+ cells represented a median of 0.5% of lymphoid cells (**Figure 5**), as compared to 50% in ITP patients and 57% in HD patients (**Figure 1**). As previously reported, PC represented the majority of residual CD19+ cells (median 73%, range 39.7 – 97.8), with memory B cells accounting for most of the other cells (25% in median, range 2.2 – 77.6). The number of residual memory B cells was highly variable among patients (median 153 cells per 10^5^ lymphoid cells, range 2 - 911), but in any case not negligible at the scale of a whole human spleen.

**Figure 5:**
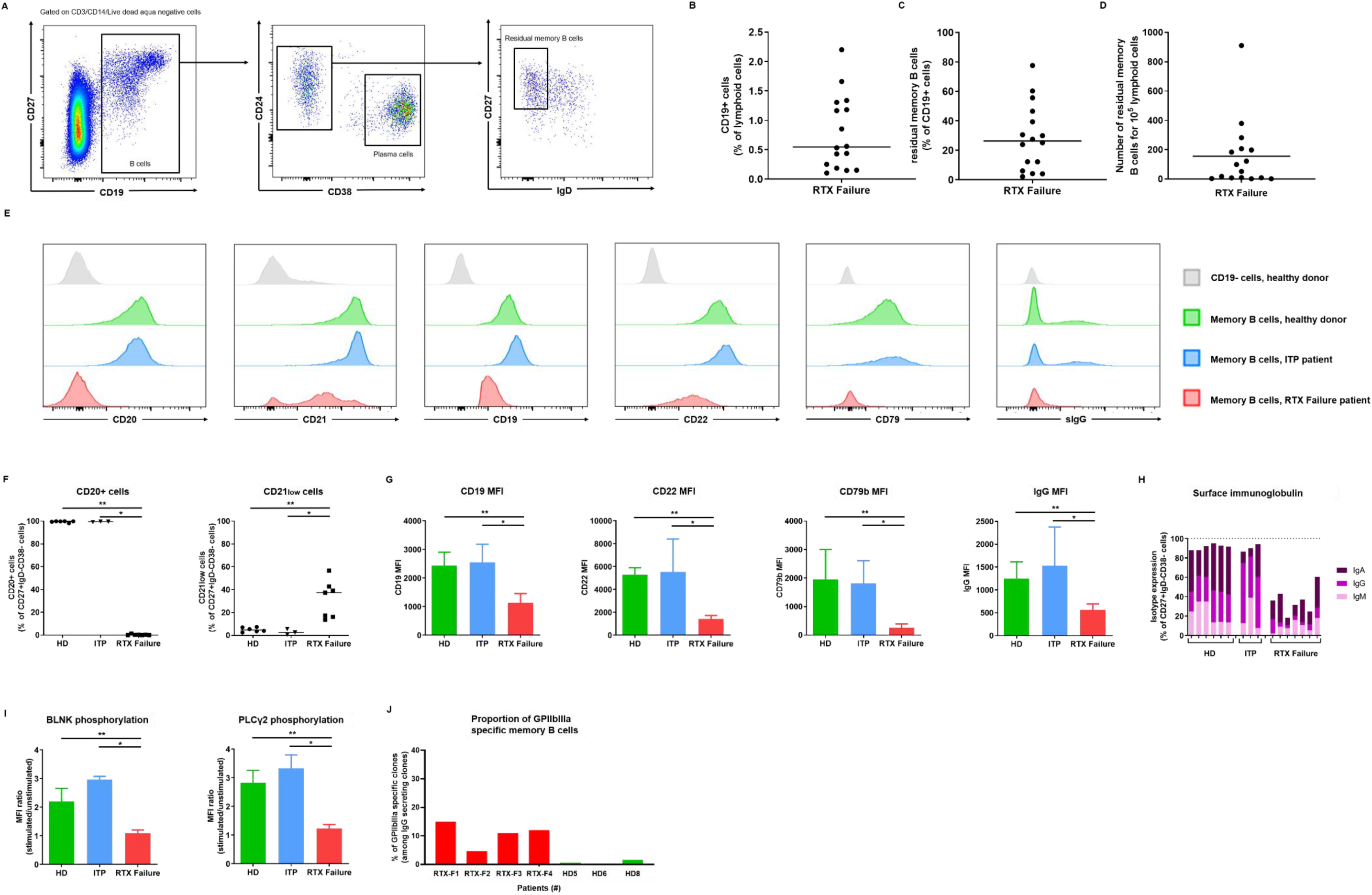
Rituximab resistant memory B cells have a unique surface phenotype and contain autoreactive clones. **(A)** Representative dot plots showing gating strategy by flow cytometry analysis of splenic mononuclear cells labeled with antibodies to CD3, CD14, CD19, CD24, CD27, CD38, IgD. After gating on CD19+ B cells among live CD3-CD14-lymphoid cells. Memory B cells were identified as CD24+, CD38-, IgD-, PC were identified as CD27+CD24–CD38^hi^. (**B-D**) Proportion of CD19+ B cells **(B)**, residual memory B cells **(C)**, and number of residual memory B cells **(D)** in the spleen of RTX failure patients (n=16). (**E**) Representative overlays of surface markers assessed by flow cytometry expressed on CD19- cells from HD (grey), memory B cells from HD (green), memory B cells from ITP patients (blue) and residual memory B cells from RTX failure patients (red). **(F)** Proportions of CD20+ cells and CD21low cells in HD (n=6), ITP (n=3), RTX failure (n=7) patients. **(G)** Histograms showing MFIs (+/-SD) of CD19, CD22, CD79b and IgG in HD (n=6), ITP (n=3) and RTX failure (n=7) patients. **(H)** Cumulative proportions of surface IgM, IgG and IgA in HD, ITP and RTX failure patients. Each bar represents one patient. **(I)** BLNK and PLCγ2 phosphorylation assessed by stimulating total splenocytes from HD, ITP and RTX failure patients with anti-Pan Ig antibodies. MFI ratio of stimulated/non-stimulated cells are indicated for each group. **(J)** Frequency of GPIIbIIIa specific clones among memory B cells in RTX failure patients (red) and healthy donors (green). The total number of tested cells is given in supplementary Table 2. Two-tailed Mann-Whitney tests were performed (\*\*\**P* < 0.001; \*\**P* < 0.01, \**P* < 0.05). Symbols indicate individual samples; horizontal bars represent median values.

Next, we focused our phenotypic analysis on the 7 RTX failure patients with higher numbers of residual memory B cells. None of these RTX-resistant cells had detectable expression of CD20 (**Figure 5**), even after intracellular staining (data not shown). ScRNAseq analysis confirmed that MS4A1 transcripts were present, albeit at a lower level compared to memory B cells from HD or ITP patients (data not shown). Unexpectedly, RTX-resistant memory B cells also had decreased surface expression of many BCR-associated molecules such as CD79b, CD19, CD21, CD22 and CD72 (**Figure 5 and Supplementary Figure 6**). Surface and intracellular immunoglobulin expression (including IgM, IgA and IgG) was also strongly reduced, resulting in a majority of cells without detectable isotype.

We assessed the expression of several surface markers on residual memory B cells known to be associated with “age-associated B-cells” (ABC; Phalke and Marrack, 2018; Portugal et al., 2017). CD11c was slightly more expressed on residual memory B cells than in memory B cells from controls, and we could also detect a minor population of residual memory B cells expressing FcRL5, but not FcRL4 (**Supplementary Figure 6**). However, we observed a reduced expression of CD19, CD72, and CXCR5 on residual memory B cells, while these markers were described to be elevated in ABC. Moreover, preliminary data obtained by scRNAseq analysis identified a subset of ABC in HD/ITP patients that clustered independently from residual memory B cells (data not shown), indicating that both populations were clearly distinct.

Residual memory B cells had limited surface expression of BCR, which is required for antigen dependent B cell activation. To confirm that these cells could not be stimulated through antigen recognition, we assessed the phosphorylation upon BCR ligation of downstream proteins BLNK and PLCγ2 involved in BCR signaling. While residual memory B cells had similar basal phosphorylation of both proteins (data not shown), BCR ligation induced BLNK and PLCγ2 phosphorylation only in memory B cells from controls (**Figure 5**). Next, we tried to stimulate residual memory B cells through BCR independent signals such as TLR and CD40L. *In vitro* stimulation by CpG and CD40L expressing cells both induced proliferation and plasma cell differentiation of residual memory B cells (**Supplementary Figure 6C**).

Finally, we searched for anti-GPIIbIIIa clones among residual memory B cells by using our culture assay described in Figure 3. This analysis showed that RTX failure patients had 4.7 to 15% of residual memory B cells reacting against GPIIbIIIa (**Figure 5**), confirming that some autoreactive clones escaped RTX depletion. Overall, these data suggest that during B cell depletion, a residual population of CD20-memory B cells that contains autoreactive clones persists but cannot be activated upon antigen encounter due to the downregulation of surface BCR expression.

Preliminary data from scRNAseq analyses identified residual memory B-cells as a unique population (data not shown), suggesting that their particular phenotype was induced by B-cell depletion rather than selected from a preexisting subset. To test this hypothesis, we cultured fresh spleen tissues from HD with and without RTX. We observed that CD20 expression was undetectable on B cells by FACS analysis as soon as 30min after incubation with RTX (data not shown). After overnight incubation with 1 and 10 µg/ml RTX, 32 and 30% of B cells were depleted in median, respectively (**Figure 6**). We also observed phenotype modifications on B cells remaining after overnight culture in presence of RTX, including reduced expression of CD19, CD79b, CD21 and CD22, but not in B cells cultured without RTX (**Figure 6**). Surface IgD and IgM expression were also reduced on CD27- B cells when cultured with RTX. These results suggested that RTX induces phenotype modifications *per se*, rather than selecting preexisting CD20 negative B cells. In agreement with this hypothesis, we did not find any CD20-CD19lowCD79lowCD21low B cells in the spleen of HD or ITP patients.

**Figure 6:**
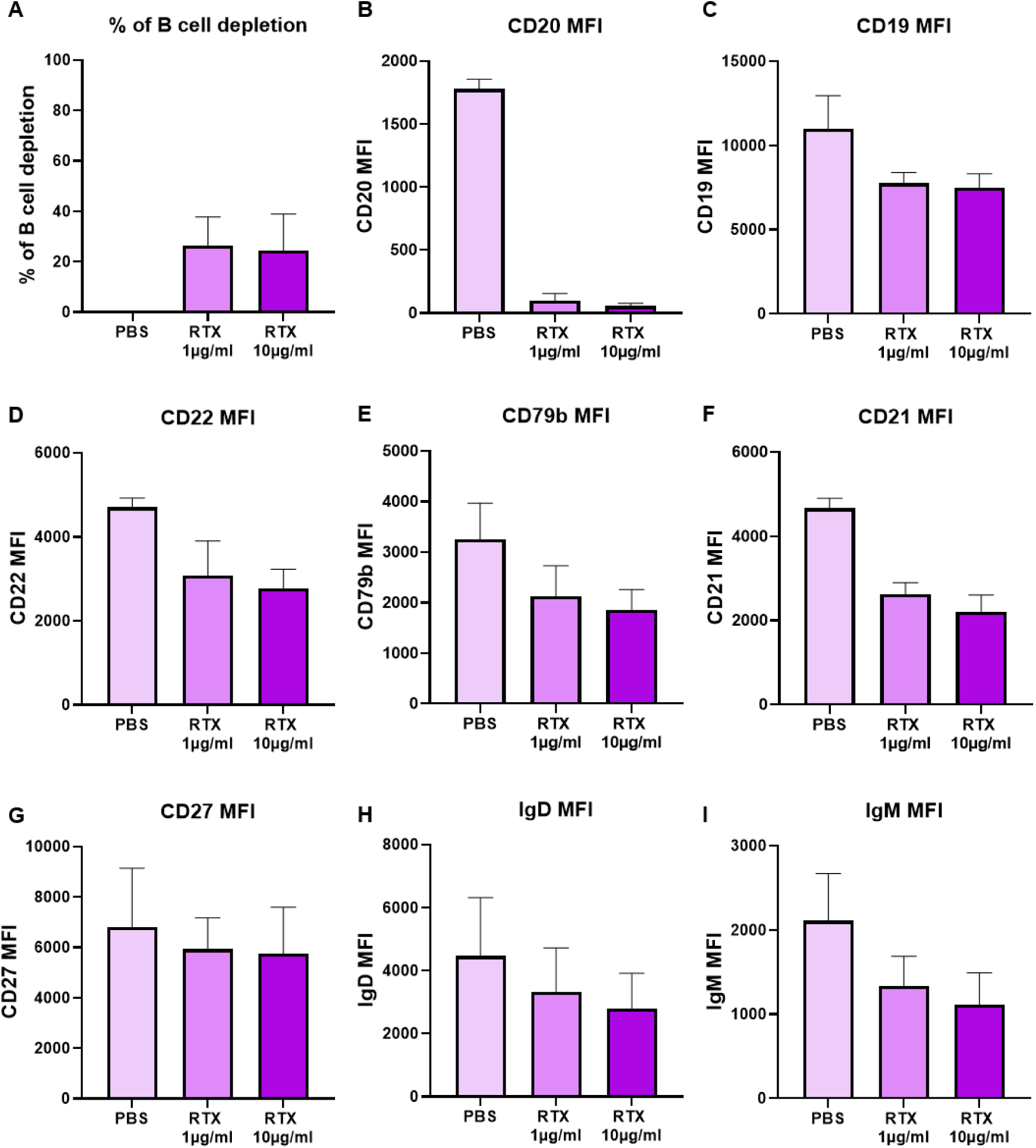
Residual memory B cell phenotype can be reproduced by in vitro exposition to rituximab. Fresh total splenocytes from HD (n=3) were incubated overnight with RTX (1 or 10 µg/ml) or PBS. **(A)** Percentage of B cell depletion, calculated with the following formula: 100 –[(B cell:T cell ratio in sample with RTX)/(B cell:T cell ratio in sample with PBS) x 100]. **(B-G)** Histograms showing MFIs (+/-SD) of CD20, CD19, CD22, CD79b, CD21 and CD27 on total B cells. **(H-I)** Histograms showing MFIs (+/-SD) of IgD and IgM on CD27- B cells. Two-tailed Mann-Whitney tests were performed (\*\*\**P* < 0.001; \*\**P* < 0.01, \**P* < 0.05).

## Discussion

RTX-resistant memory B cells have been described in lymphoid organs (Audia et al., 2011; Kamburova et al., 2013; Mahévas et al., 2012; Ramos et al., 2007; Ramwadhdoebe et al., 2019; Wallin et al., 2014), but their exact contribution to disease relapses has never been formally addressed. Autoreactive PC could arise from these residual memory B cells, and/or from autoreactive clones contained in the naïve repertoire generated during B cell reconstitution. Spleens of ITP patients relapsing after an initial response to B-cell depletion offer a unique opportunity to study the precise onset of disease relapses and explore these different scenarios.

In RTX relapse patients, GC were significantly and largely expanded as observed in ITP patients, suggesting that they were central in disease reinitiation. This contrasted with a severely reduced memory B cell population that nonetheless contained proliferating cells. To better understand the relationships between GC B cells and other populations, we performed high throughput sequencing of immunoglobulin heavy chain genes of GC B cells, memory B cells and PC. Because mutation accumulation in GC is a time-dependent process (Reynaud et al., 2012), we used the number of mutations in VH segment as a surrogate marker to estimate B cell resistance to RTX. This was clearly exemplified in the PC, that are known to resist RTX and displayed a bimodal distribution of somatic mutations contrasting with the gaussian distribution expected in this populations (Bagnara et al., 2015; Tipton et al., 2015). Strikingly, analysis of clonal relationships supervised by a cut-off value selected according to the bimodal distribution confirmed that low mutated and highly mutated cells originated from distinct populations. This revealed the coexistence of two synchronous responses in GC of RTX relapse patients, coming from RTX-resistant and newly generated B cells.

We developed a single cell culture assay to identify and sequence B-cell clones reactive against GPIIbIIIa, the main target antigen in ITP (He et al., 1994). Although we cannot exclude that cross-reactive or polyreactive clones could be detected with this system, as suggested by the presence of a few anti-GPIIbIIIa clones in GC from HD, there was a clear enrichment in anti-GPIIbIIIa GC and memory B cells from RTX relapse patients. This contrasted with the relatively low frequency of anti-GPIIbIIIa PC, in agreement with our previous findings (Mahévas et al., 2012). This discrepancy could suggest a mechanism of regulation preventing autoreactive GC B cells to differentiate into PC. More probably, PC generated from GC in RTX relapse patients are diluted in the whole PC population that resisted RTX, whereas GC and memory B cell populations were almost completely depleted by anti-CD20 and are thus enriched in autoreactive cells during relapses. We cannot exclude that autoreactive PC differentiated into short-lived plasmablasts that either migrated to the peripheral blood or were damped by steroid therapy. Because anti-GPIIbIIIa IgM ELISPOT on splenocytes is an unreliable assay, we could not estimate the frequency of anti-GPIIbIIIa IgM secreting PC that may represent the major part of the new autoimmune response.

Analysis of VH sequences from anti-GpIIbIIIa clones indicated that some memory B cells were highly mutated, probably corresponding to RTX-resistant cells. Such cells were present in all patients with detectable anti-GPIIbIIIa B cells, indicating that it was not a rare event. In contrast, the majority of anti-GpIIbIIIa GC B cells harbored less than 10 mutations, although more mutated clones were clearly identified in some patients. This reflected the predominance of newly generated B cells in GC, in accordance with the high proportion of IgM isotype in GC. We found that some highly mutated anti-GPIIbIIIa clones from the 3 patients were present in GC, memory and PC populations, confirming that RTX resistant memory B cells directly participate in ITP relapses by joining GC reaction and/or differentiating into autoreactive PC. Clonal tree reconstruction confirmed the position of memory B cells as ancestors of either PC or GC B cells. We also observed low mutated anti-GPIIbIIIa clones shared between GC and PC populations, confirming that newly generated B cells contributed to disease relapses as well. Why and how naïve B cells are recruited in autoreactive GC remains to be assessed, but overall, our results suggest that GC reactions are reinitiated by RTX-resistant autoreactive memory B cells. Indeed, memory B cells can activate more quickly and more efficiently than naïve B cells in response to antigen stimulation in secondary responses. It has also been shown in a transgenic mouse model that a single autoreactive B cell clone can initiate a germinal center reaction where wild-type B cells acquiring an autoreactive specificity rapidly predominate (Degn et al., 2017). T-follicular helper cells (TFH) are central in GC formation and maintenance (Ueno et al., 2015) and have been shown to be dysregulated in ITP (Audia et al., 2014). Even if it remains speculative in our setting, memory B cells are efficient antigen presenting cells and could reactivate cognate TFH, thus favoring disease reinitiation (Asrir et al., 2017; Ise et al., 2014).

Several studies have documented the presence of residual B-cells in secondary lymphoid organs after RTX, with variable proportion of memory B cells (Audia et al., 2011; Kamburova et al., 2013; Mahévas et al., 2012; Ramos et al., 2007; Ramwadhdoebe et al., 2019; Wallin et al., 2014). In RTX failure patients, residual memory B cells did not express surface CD20, contrasting with the presence of *MS4A1* transcripts. Several mechanisms accounting for the lack of CD20 detection have been extensively described *in vitro* and *in vivo*, such as epitope masking by RTX, antigen modulation through CD20 internalization upon RTX binding (Lim et al., 2011; Reddy et al., 2015; Vaughan et al., 2014) or trogocytosis (Beum et al., 2011; Boross et al., 2012; Rossi et al., 2013; Taylor and Lindorfer, 2015). Interestingly, we found a strong reduction of BCR expression, including surface and intracellular immunoglobulin and CD79b molecule, on residual memory B cells. Accordingly, residual memory B cells did not respond to BCR stimulation but could still differentiate into plasma cells *in vitro* upon CD40 or TLR stimulation. Type I anti-CD20 antibodies such as RTX are known to induce CD20 aggregation and migration to lipid rafts (Boross and Leusen, 2012). Additionally, they induce a colocalization of the BCR and CD20 molecules on target cells, resulting in a calcium influx triggered by BCR signaling (Walshe et al., 2008). Most BCR co receptors (including CD19, CD22, CD72) were downmodulated on residual memory B cells. Despite their low or absent CD21 expression, CD20-memory B cells differ phenotypically from age-associated B cells (also termed CD21low B cells or atypical memory B cells). Whether lack of BCR coreceptors is directly linked with the BCR absence itself or the indirect consequence of trogocytosis as previously suggested remains an open question (Rossi et al., 2013; Taylor and Lindorfer, 2015). This unique phenotype could explain why residual memory B cells remain inactive, despite their autoreactivity potential, until RTX has been cleared.

RTX has been shown to inhibit anti-apoptotic pathways *in vitro* (Bonavida, 2007). It is therefore surprising to observe that some residual memory B cells survived for months *in vivo* despite constant anti-CD20 exposure. Analysis of the molecular program of these cells will probably provide a part of the answer. We can speculate that RTX, by modifying the microenvironment, enable the emergence of a particular transcriptional program in memory B cells favoring their survival, as previously described for LLPC (Mahévas et al., 2012, 2015; Thai et al., 2018).

From a clinical perspective, targeting residual memory B cells could represent an attractive therapeutic option to avoid relapses. Several strategies could be used, such as anti-CD19 antibodies which are currently in development in autoimmune diseases and could also limit the emergence of long-lived PC observed after RTX under the influence of the microenvironment. Optimizing B cell depletion with the use of new-generation anti-CD20 such as obinutuzumab could also limit the persistence of memory B cells. Because analysis of secondary lymphoid organs after RTX administration is limited in humans, it remains to be demonstrated whether our findings can be generalized to other autoimmune diseases where RTX is effective. However, as mentioned before, RTX-resistant B cell populations mainly composed of memory B cells have been described in several auto-immunes diseases including RA (Ramwadhdoebe et al., 2019) and AIHA (Mahévas et al., 2015), as well as after preventive treatments of humoral graft rejection (Kamburova et al., 2013; Ramos et al., 2007; Wallin et al., 2014), suggesting that persistence of autoreactive clones is not limited to ITP.

Collectively, our results strongly suggest that splenic autoreactive residual memory B cells survived for months after RTX administration and actively participate in disease relapses during B cell reconstitution, through the reinitiation of the autoreactive GC response, which was further amplified by newly generated B cells. Our study paves the way for the development of new strategies to optimize B cell depletion in order to increase the rate of long-term remissions in antibody-mediated diseases.

## Methods

### Patients

#### ITP patients

All patients included in this study were adults, and had ITP diagnosis according to the international guidelines definitions (Rodeghiero et al., 2009). Patients with underlying immunodeficiency, hepatitis C virus infection, lymphoproliferative disorders, thyroid or liver disease, and defined systemic lupus erythematosus (≥ 4 American Rheumatism Association criteria) were excluded. All patients had persistent (≥ 3 months) or chronic (≥ 12 months) ITP. Complete response was defined as platelet count >100×109/L and no bleeding, and failure as platelet count <30×109/L (Rodeghiero et al., 2009).

#### Control patients

All control patients were organ donors that died from stroke. None of them had lymphoma or autoimmune disease.

#### Spleen sample processing

Immediately after splenectomy, mononuclear cells were isolated from crushed pieces of splenic tissue by ficoll density-gradient centrifugation and stored in liquid nitrogen.

### Histology

Fixed tissue samples from spleen of RTX relapse patients were investigated after Hematoxylin and eosin staining (HES) and immunohistochemical stainings including CD20 (M755-01, DAKO), CD79a (M705001, Agilent), CD3 (A0452, DAKO), Ki67 (M7240, DAKO), CD10 (CD10-270-L-CE, Leica), CD21 (CD21-2G9-L-CE, Leica), and PD1 (ab52587, Abcam).

### Multi-color flow cytometry and cell sorting

After thawing, mononuclear cells were counted and stained with antibodies directed against surface markers (listed in **Supplementary Table 5**). Intracellular stainings were performed after using BD Cytofix/Cytoperm solution (BD Biosciences). Stained populations were analyzed on a FACS Fortessa LSR with DIVA (BD Biosciences) and FlowJo (Treestar Inc.) softwares. Cell sorting was performed with a FACS Aria IIIu cell sorter (BD Biosciences).

### Single cell culture

Single cell culture was performed as previously described (McCarthy et al., 2018). Single B cells were sorted in individual wells containing MS40 cells expressing CD40L (gift from G. Kelsoe). Cells were cocultured at 37°C with 5% CO2 during 15 days in RPMI-1640 (Invitrogen) supplemented with 10% HyClone FBS (Thermo Scientific), 55 µM 2-mercaptoethanol, 10 mM HEPES, 1 mM sodium pyruvate, 100 units/mL penicillin, 100 mg/mL streptomycin, and MEM non-essential amino acids (all Invitrogen), with the addition of recombinant human BAFF (10 ng/ml), IL2 (50 ng/ml), IL4 (10 ng/ml), and IL21 (10 ng/ml; all Peprotech). Half of supernatant was carefully removed at days 4, 8 and 12 and the same amount of fresh medium with cytokines was added to the cultures. IgG ELISA was performed at day 15 for all wells to screen for IgG secreting clones.

### ELISPOT assays

Anti-GPIIbIIIa ELISPOTS were performed as previously described (Mahévas et al., 2012). Briefly, 2.5 μg/ml keyhole limpet hemocyanin (KLH), 10 μg/ml goat anti-human Ig polyvalent antibody (Invitrogen), or 15 μg/ml purified GpIIbIIIa (Stago) were coated in PBS/CaCl2 0.05% in multiscreen 96-well filter plates (MSIPS4510; Millipore) with overnight incubation at 4°C.

#### Identification of anti-GPIIbIIIa specific cultured clones

IgG secreting clones from single cell cultures of memory B cells or germinal center B cells were transferred in the ELISPOT plate in individual wells and incubated overnight at 37°C with 5% CO2. Cells were then carefully removed and stored at 37°C, and the ELISPOT plate was incubated with biotinylated goat anti-human IgG Fc (Invitrogen), followed by HRP-conjugated avidin (Vector Laboratories) and developed using 3-Amino-9-ethylcarbazole (BD biosciences). Spots were enumerated in each well with an ELISPOT reader using the AID software (version 3.5; AutoImmun Diagnostika). Clones were considered GPIIbIIIa reactive if more than 3 dense and specific spots were detected in the well. AP3 hybridoma was purchased from ATCC.

#### Identification of anti-GPIIbIIIa specific splenic plasma cells

Alternatively, 10^6^ splenocytes were serially diluted in culture medium in triplicate before transfer to ELISPOT plates and incubated overnight at 37°C with 5% CO2. IgG-secreting anti-GpIIbIIIa spots were enumerated in each well and reported to IgG-positive spots.

### IgH sequencing of GPIIbIIIa specific clones

GPIIbIIIa specific clones identified in the ELISPOT assay were washed in PBS, and RNA was extracted using RNeasy micro kit (Qiagen) according to the manufacturer protocol. Reverse transcription was performed using SuperScript III enzyme (ThermoFisher) in 12 μl volume (42°C 10min, 25°C 10min, 50°C 60min, 94°C 5min) with 6 µl of RNA and random hexamers (GE Healthcare). A PCR was then performed based on an established protocol (Tiller et al., 2008). Briefly, 3.5 μl of cDNA were used as template and amplified in a total volume of 40 μl with forward L-VH primer mix and reverse Cγ primer (sequences in **Supplementary Table 6**), using HotStar® Taq DNA polymerase (Qiagen) for 30 cycles (94°C 30s, 58°C 30s, 72°C 55s). PCR products were sequenced with the reverse primer CHG-D1 and read on ABI PRISM 3130XL genetic analyser (Applied Biosystems). Sequence quality was verified with the CodonCode Aligner software (CodonCode Corporation), and data were analyzed with the IMGT/HighV-QUEST web portal (from The International Immunogenetics Information System).

### Recombinant antibody production and purification

IgH and IgL chains from clones of interest were cloned as previously described (Tiller et al., 2008). HEK293T cells were transfected with plasmid DNA encoding paired IgH and IgL chains using Jet Prime kit (Polyplus) as previously described (Tiller et al., 2008). Recombinant antibodies were purified from supernatants using protein-G beads (GE Healthcare).

### ELISA and MAIPA

Total IgG and anti-GPIIbIIIa IgG from culture supernatants were detected by home-made ELISA. Briefly, 96 well ELISA plates (Thermo Fisher) were coated with either goat anti-human Ig (10μg/ml, Invitrogen) or purified GPIIbIIIa (5µg/ml, Stago) in sodium carbonate. Cell culture supernatants were added for 1h, then ELISA were developed using HRP-goat anti-human IgG (1μg/ml, Immunotech) and TMB substrate (Eurobio). OD450 and OD620 were measured and Ab-reactivity was calculated after subtraction of blank wells.

Indirect MAIPA assay (Apdia) was performed following the manufacturer instructions.

### Proliferation assay

Cell proliferation was assessed with Cell proliferation Dye eFluor450 (Thermo Fisher). Briefly, splenocytes were resuspended at 2.5×10^6^ cell/ml in BCM and stained according to the manufacturer instructions. Stained cells were cultured for 5 days in BCM supplemented or not with CpG (ODN2006, 2,5µg/ml, Invivogen), or cocultured with MS40 cells expressing CD40L and cytokine cocktail (recombinant human BAFF (10 ng/ml), IL2 (50 ng/ml), IL4 (10 ng/ml), and IL21 (10 ng/ml), all Peprotech). Cell division was assessed by measuring the decrease in Dye eFluor450 fluorescence via flow cytometry on memory B cells (CD3-CD14-CD19+CD38-CD24+CD27+IgD-).

### In vitro RTX cultures

Fresh spleen samples from healthy donors were obtained and immediately crushed. Total splenocytes were seeded at 5×10^6^ cells/well in 24-well plates with complete B cell medium. RTX (1µg/ml or 10µg/ml) or PBS was added to the culture. After overnight incubation at 37°C with 5% CO2, cells were washed and stained for FACS analysis.

### Library Preparation and Illumina Sequencing

Splenic B cell subsets from three RTX relapse patients, including PC (CD19+CD27highCD38highCD24-), GC (CD19+CD38intCD24-IgD-CD20+), memory (CD19+CD38-CD24+CD27+IgD-) and naïve (CD19+CD38-CD27-IgD+) B cells, were sorted in PBS/2% FCS. 10^4^ to 2×10^5^ cells were obtained for each subset. Cells were washed with PBS, centrifugated, and resuspended in the appropriate volume of Lysis/Binding Buffer from the Dynabeads mRNA DIRECT Micro Kit (ThermoFisher). mRNA were isolated directly with dynabeads oligo (dT) and immediately used in their entirety for reverse transcription in solid phase, according to a protocol adapted from Vergani et al (Vergani et al., 2017), with modified primers (**Supplementary Table 6**). Briefly, cDNA were synthetized in 10 μl (50°C 1 h, 72°C 10 min) using SuperScript III Enzyme (ThermoFisher). After RNase H treatment, second-strand synthesis was performed in solid phase in 10 μl using Q5 Polymerase (NEB) and a mix of 13 primers covering IGHV leader sequence, containing 13 to 16 random nt and partial Illumina adaptor sequences (37°C 20 min, 98°C 30 s, 62°C 2 min, and 72°C 10 min). IGHV-D-J-CH double-stranded cDNA attached to the magnetic dynabeads were washed three times in 10 mM Tris–HCl pH 8.0 to remove the remaining primers, and the entire sample was used as template for the PCR1 amplification in 10 μl using Q5 Polymerase with a forward primer that is specific for the partial Illumina adaptor sequence and allowing its extension, and a mix of reverse isotype specific (mu, alpha, gamma) primers (98°C 30 s; 10 cycles of 98°C 10 s, 58°C 15 s, and 72°C 1 min; 72°C 10 min). Depending on the initial cell fraction, individual semi-nested PCR2 were performed with inner reverse primers specific for the mu, alpha, and gamma constant region and allowing the introduction of partial Illumina adaptors. Two µl of the PCR1 product were used for the semi-nested PCR2 reactions carried in 20 μl (98°C 30 s; 15 cycles of 98°C 10 s, 58°C 15 s, and 72°C 1 min; 72°C 10 min). The PCR2 products were purified with Ampure XP beads at a ratio of 1:1, and 1–10 ng were used to add Illumina Index with the Nextera XT kit (Illumina). After a final purification with Ampure XP beads (ratio 1:1), the library obtained for each sample was diluted at a concentration of 2nM, and an equimolar pool of all the libraries was sequenced on the MiSeq Illumina with the V3 kit (2 × 300) after being loaded at 12 pm with 10% PhiX.

### Bioinformatics analysis of IgH sequencing data

UMI Barcoded Illumina MiSeq 2×300 reads were analysed with the packages of the Immcantation framework as well as custom Python, Bash and R scripts. Raw read preprocessing was done with pRESTO (REpertoire Sequencing TOolkit) (Vander Heiden et al., 2014). Sequences with average Phred quality scores less than 20 were excluded from further analysis. PCR primers were identified with a mismatch rate of 0.11 for C-primer and 0.2 for V-primer. UMI read groups having high error statistics were removed at 0.1 threshold, gap positions were removed if present in more than 50% of the sequences.

Following consensus build, the paired reads were assembled, deduplicated and submitted to IMGT/HighV-Quest web portal (the International ImMunoGeneTics database), for annotation, mutation analysis and functionality assessment (Alamyar et al., 2012). Annotated sequences with V-region length less than 250 or with number of detected somatic hypermutation exceeding 60 were filtered out.

Clonal clusters assignment and germline reconstruction was performed with Change-O toolkit (Gupta et al., 2015) on the sequences having at least two representative reads. The sequences that had the same V-gene, same J-gene, including ambiguous assignments, and same CDR3 length with maximal nucleotide hamming distance of 0.16 (the threshold determined with SHazaM R package) were considered belonging to the same clonal group.

Further clonal relationship and statistical analysis was implemented in R. The graphics were obtained with packages ggplot2, circlize, as well as with GraphPad tool. Phylogenetic trees were made using Alakazam package and Phylip tool.

### Statistics

Kruskal-Wallis test and Mann-Whitney test were used to compare continuous variables as appropriate. A *P*-value ≤ 0.05 was considered statistically significant. Statistical analyses involved use of GraphPad Prism 4.0 (La Jolla, CA, USA).

### Study approval

This study was conducted in compliance with the Declaration of Helsinki principles and was approved by the Agence de la Biomédecine and the Institutional Review Boards Comité de Protection des Personnes (CPP) Ile-de-France IX (for ITP patients) and II (for control patients). All ITP patient provided written informed consent before collecting splenic samples.

