## Supplementary material for "A reservoir of rituximab-resistant splenic memory B cells contributes to relapses after B-cell depletion therapy"

**A**

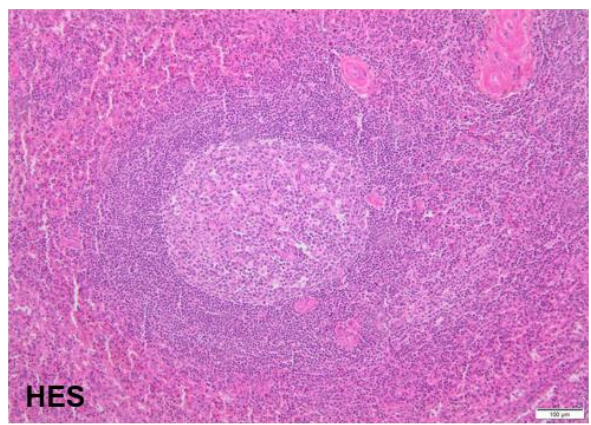

**B**

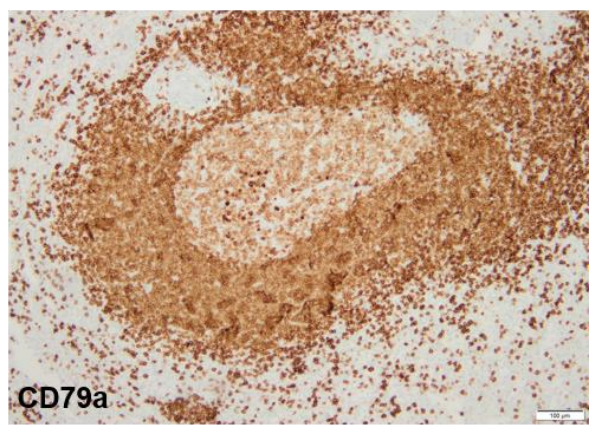

**C**

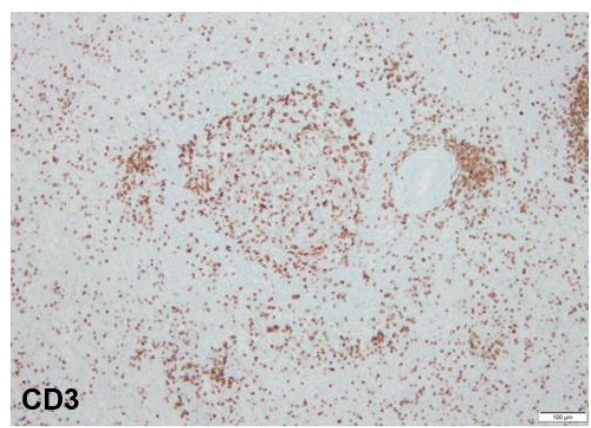

**D**

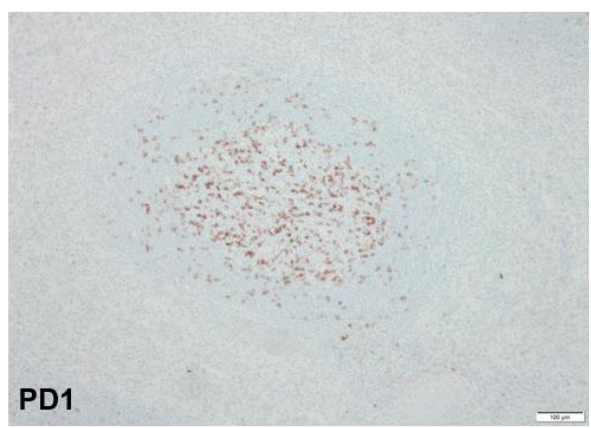

**E**

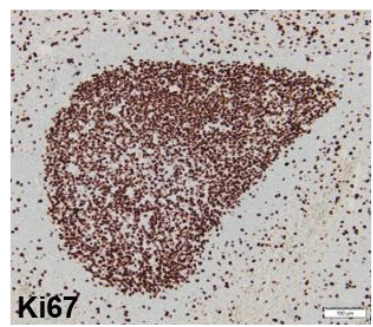

**F**

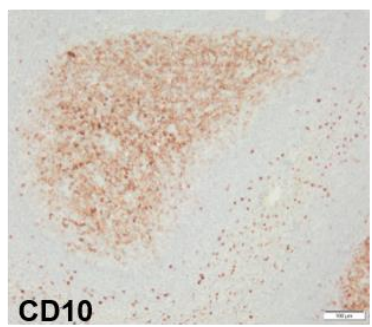

**G**

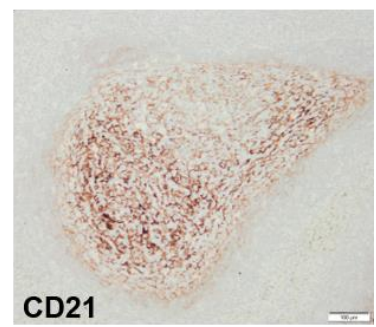

#### **Supplementary Figure 1: Germinal centers in RTX relapse patients**

Representative spleen sections from RTX relapse patients (n=2) stained with hematoxylin and eosin (**A**), CD79a (**B**), CD3 (**C**), PD-1 (**D**), Ki67 (**E**), CD10 (**F**) and CD21 (**G**) identifying GC B cells (CD79a<sup>low</sup>, CD10+, Ki67+), T Cells (CD3+), TFH cells (PD-1+), and follicular dendritic cells (CD21+). Scale bars: 100 μm.

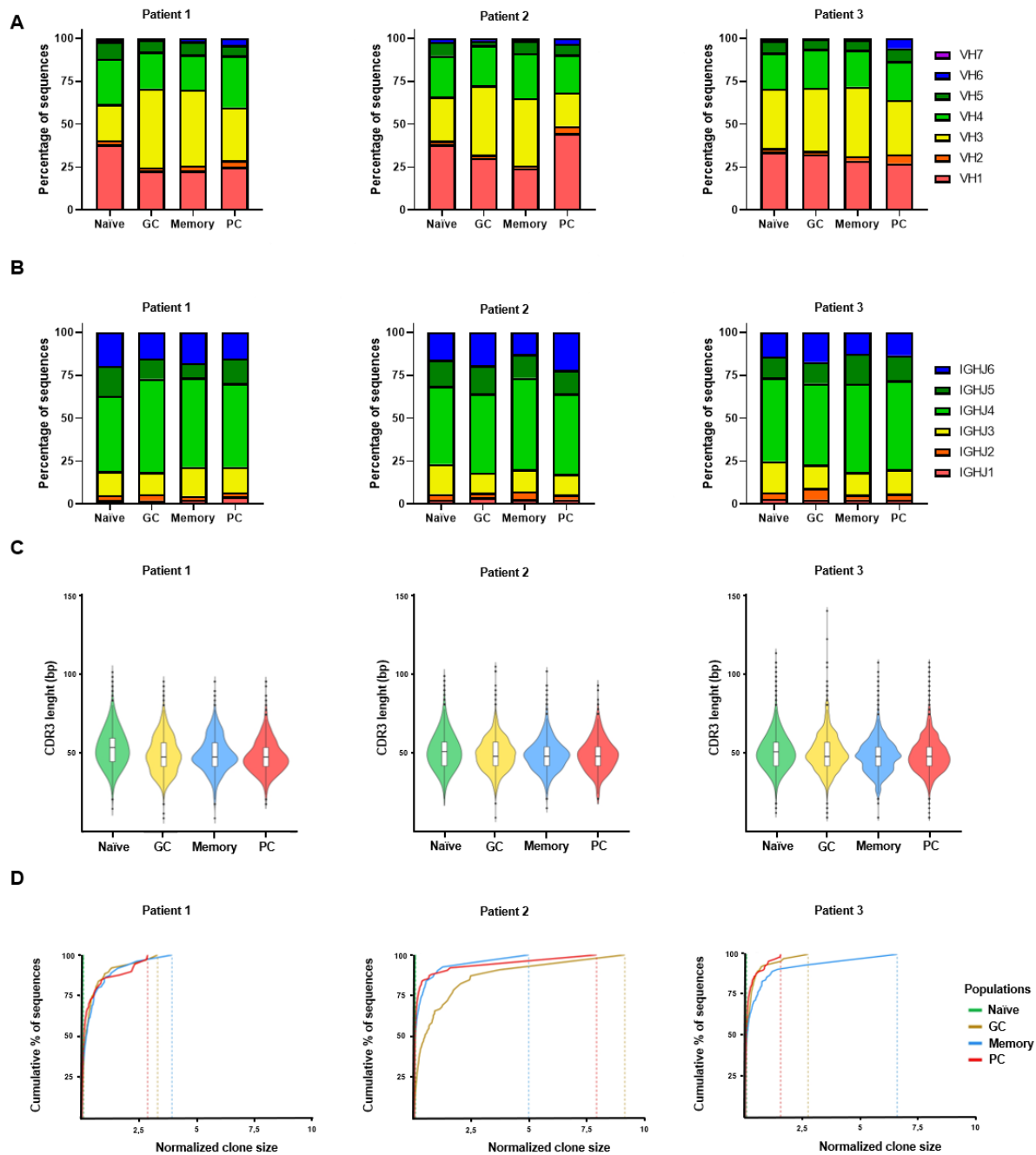

### **Supplementary Figure 2: Splenic B cell repertoire in ITP relapse patients**

High throughput IgH sequencing was used to analyze the splenic B cell repertoire of 3 RTX relapse patients. VH (A) and JH (B) usage in sequences from naïve, GC, memory and PC populations. (C) Violin plots showing CDR3 length distribution (in base pairs) in splenic B cell subsets. (D) Clonality of B cells subsets repertoire is shown by plotting the frequency of sequences in each individual clone among total sequences (clone size, horizontal axis), versus the contribution of individual clones to sequences (vertical axis) from smallest (bottom) to greatest (top). Naïve B cell repertoire (green line) containing only small clones merges with the vertical axis, while antigen-experienced B cell repertoires exhibit variable clonal expansions.

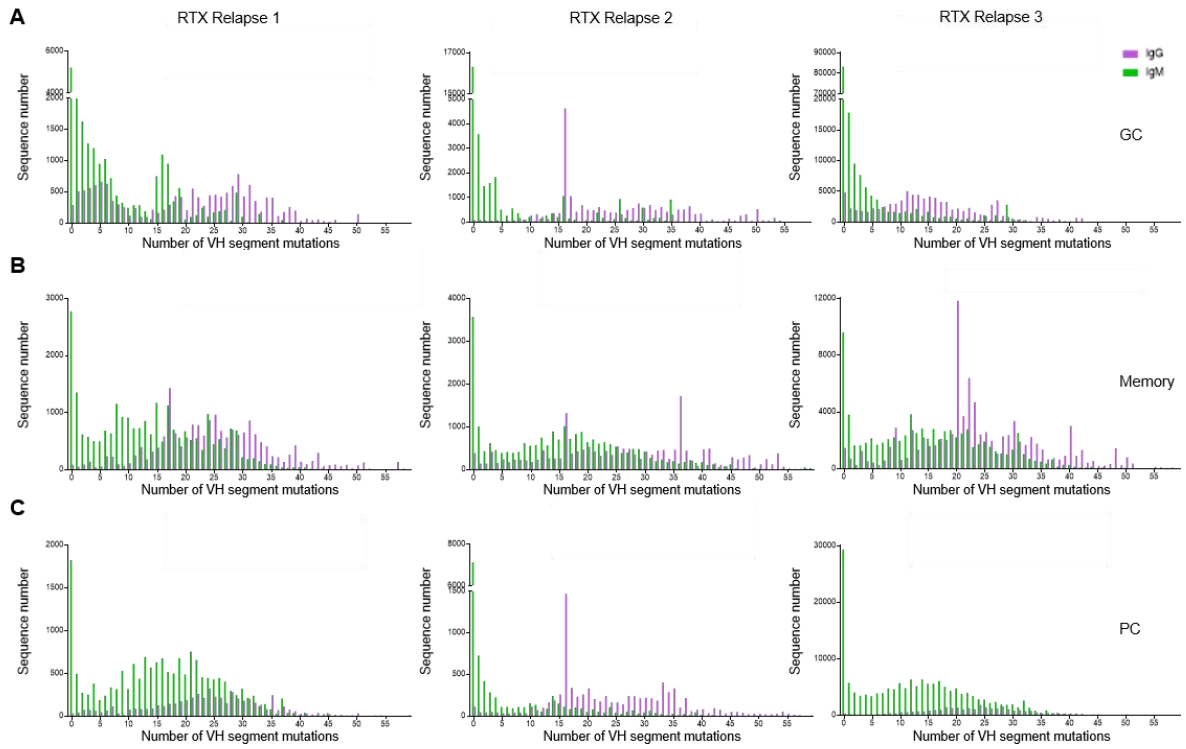

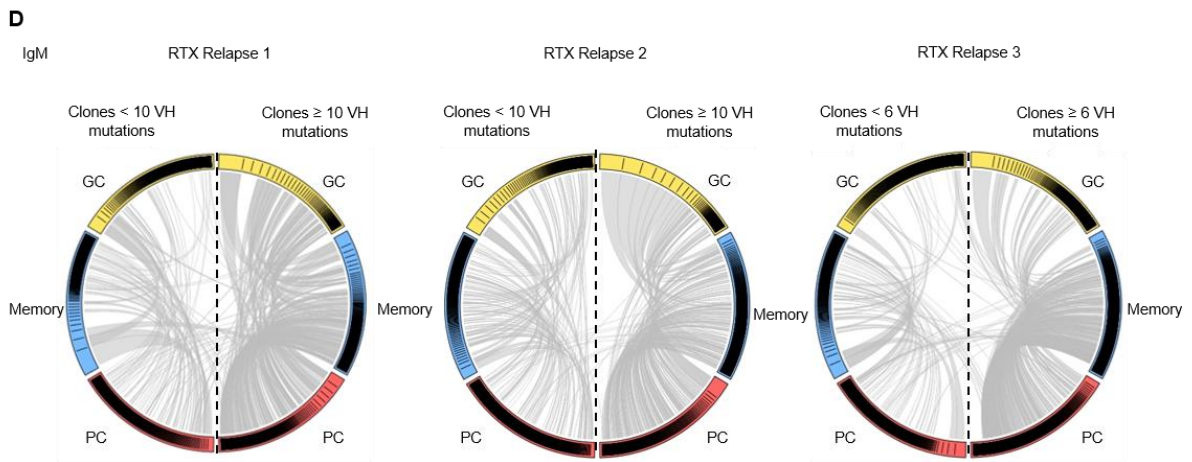

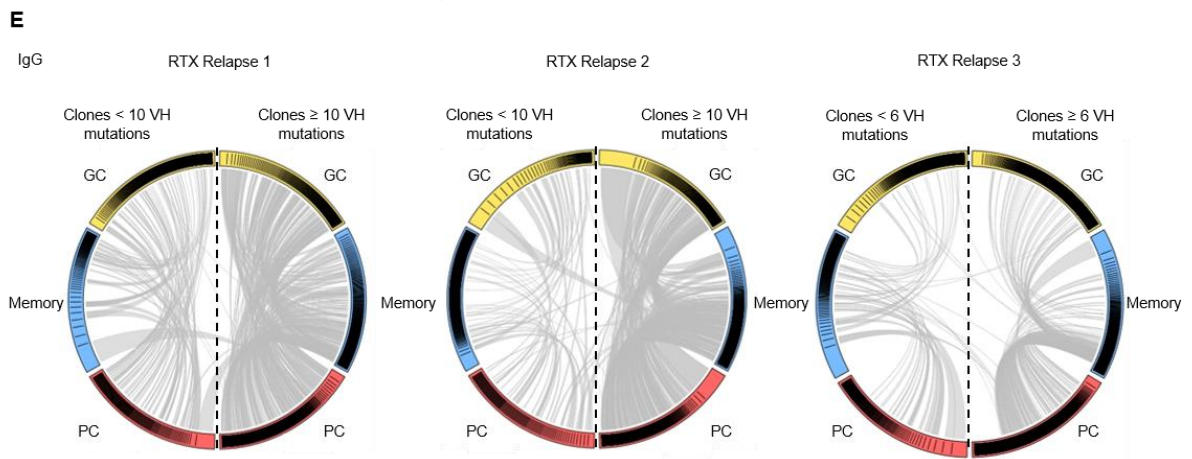

#### **Supplementary Figure 3: IgM sequences predominate in newly generated B cells and IgG sequences in rituximab resistant cells**

(A-C) VH segment mutation distribution in IgM (green) and IgG (purple) sequences from splenic GC (A), memory (B) and PC (C) populations from 3 RTX relapse patients assessed by high-throughput IgH sequencing.

Circos plot showing clonal relationships shared between IgM (D) and IgG (E) sequences from GC, memory and PC splenic populations. Clones from each population were classified into “low mutated” (left side of the plot) or “highly mutated” (right side of the plot) based on the clone median mutation number. Each colored sector represents one subpopulation and is divided into segments representing individual clones in rank-sized order. The internal connections show clones shared by multiple subpopulations.

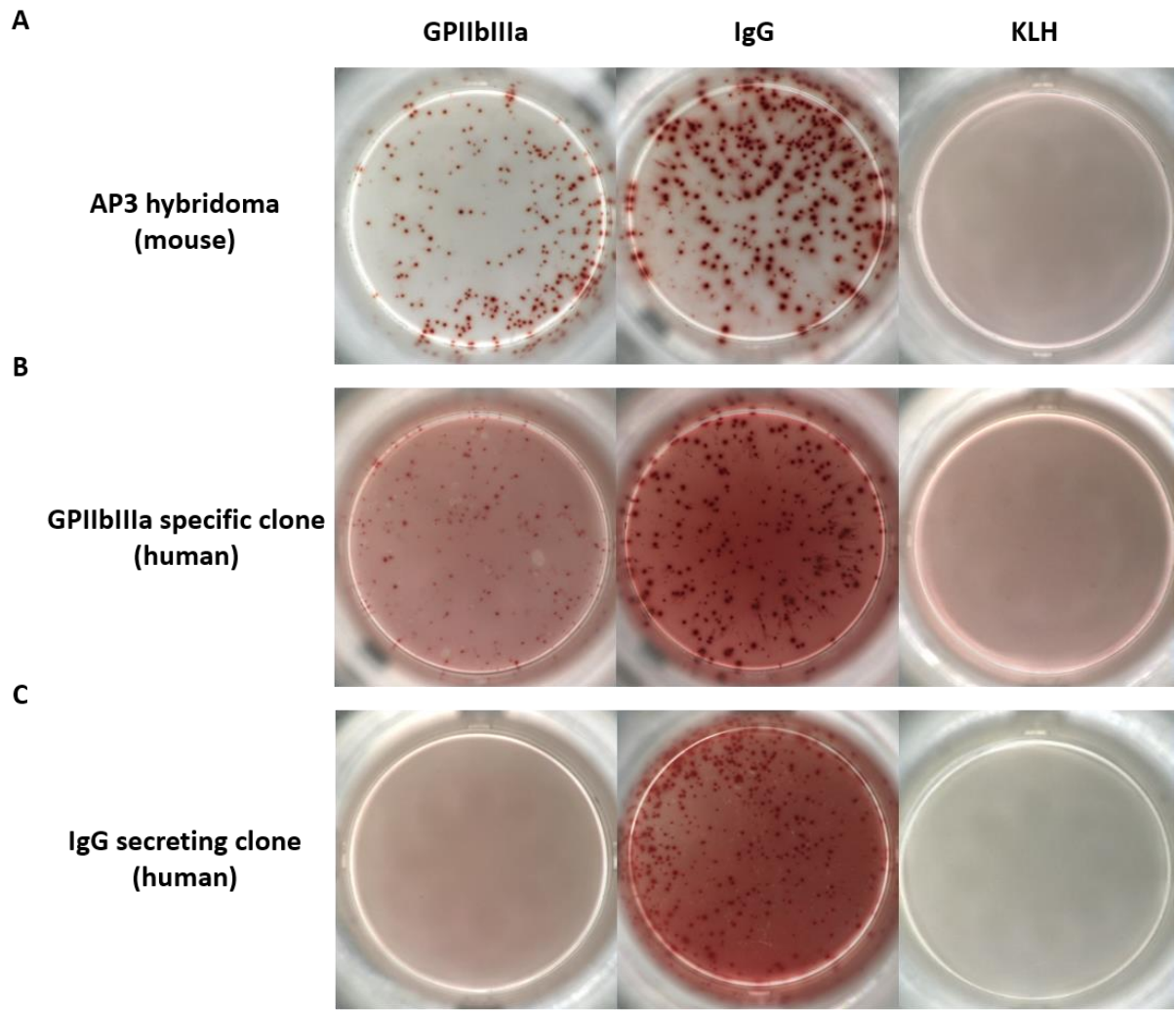

**D**

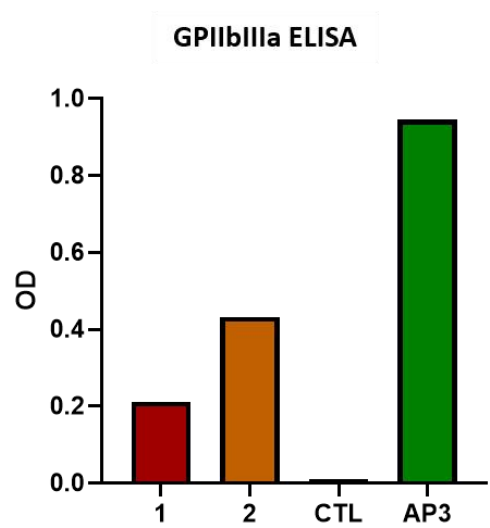

**E**

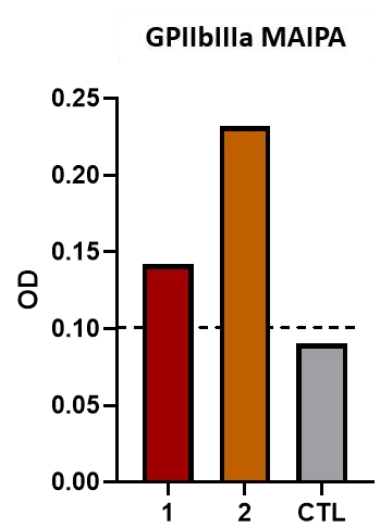

##### **Supplementary Figure 4: IgG anti-GPIIbIIIa ELISPOT allows detection of autoreactive clones**

IgG anti-GPIIbIIIa ELISPOT was used for the determination of clone specificities after single cell culture. AP3 hybridoma producing mouse IgG1 anti-GPIIbIIIa antibodies was used for the validation of IgG anti-GPIIbIIIa ELISPOT. Representative pictures of wells coated with GPIIbIIIa, anti-human immunoglobulin and Keyhole limpet hemocyanin (KLH) and tested with AP3 hybridoma (**A**), or clones from single memory B cells specific (**B**) or not (**C**) for GPIIbIIIa after 15 days of culture.

Immunoglobulin VH/VL genes from two clones (1 and 2) originating from GPIIbIIIa-specific memory B cells were sequenced and cloned into HEK cells. Anti-GPIIbIIIa ELISA (**D**) and MAIPA (**E**) performed with culture supernatants confirmed GPIIbIIIa reactivity. Irrelevant monoclonal human IgG1 was used as a negative control (CTL). Dotted line represents positive threshold for MAIPA assay.

A

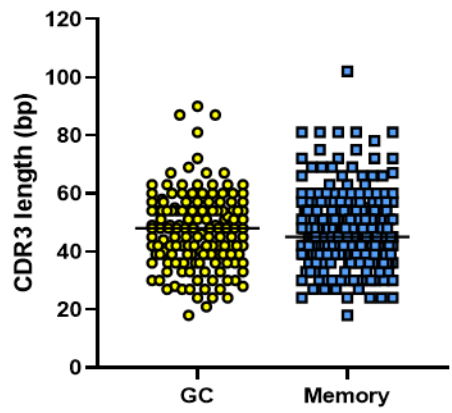

B

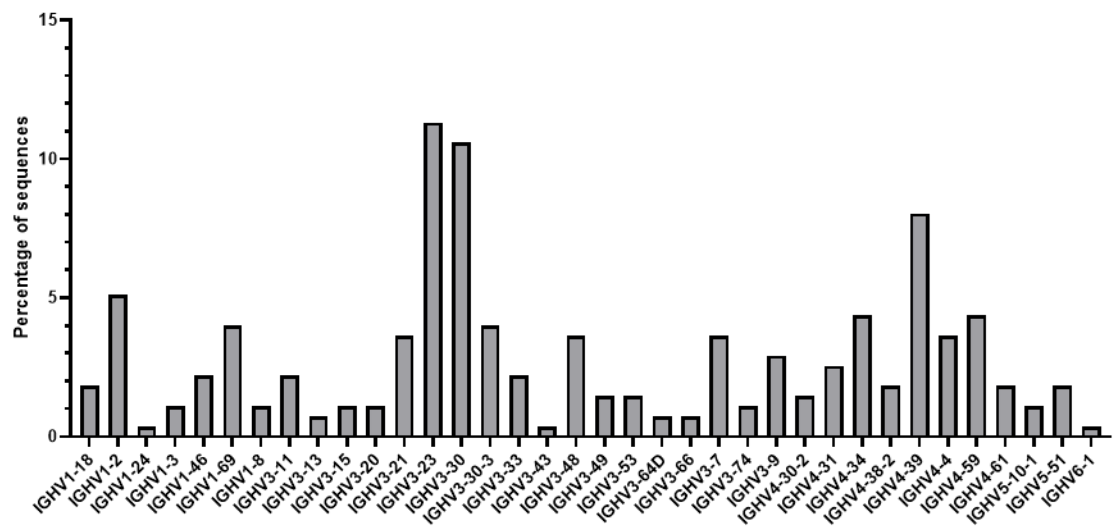

#### **Supplementary Figure 5: GPIIbIIIa specific B cells repertoire**

Sequences obtained from GPIIbIIIa specific B cells of RTX relapse patients were analyzed for CDR3 length (**A**) and VH gene distribution (**B**).

A

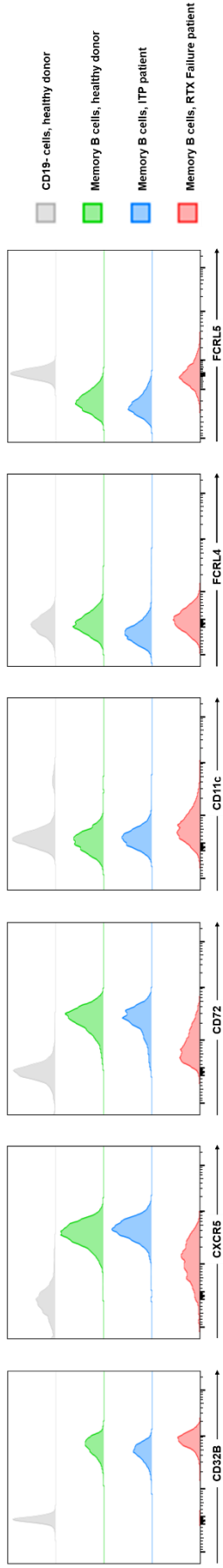

B

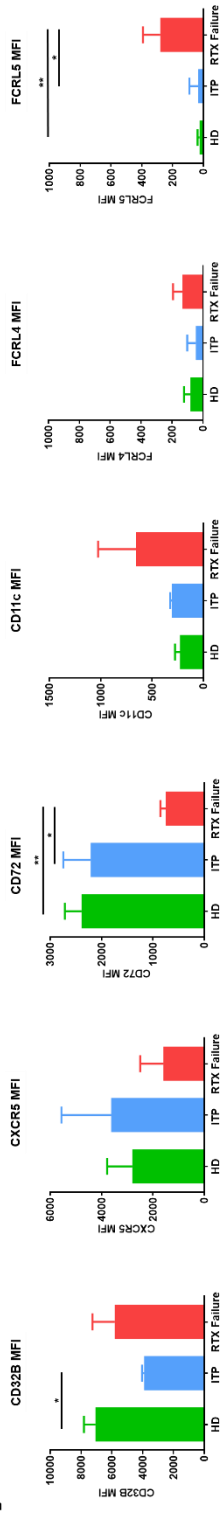

C

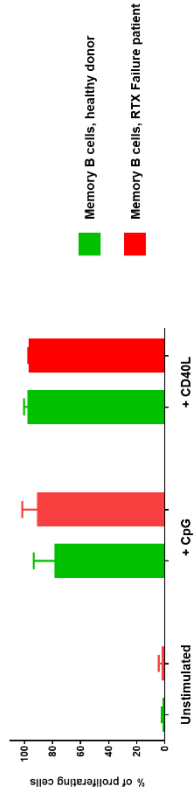

#### **Supplementary Figure 6: Rituximab resistant memory B cells can be activated in vitro**

**(A)** Representative overlays of surface markers assessed by flow cytometry expressed on CD19<sup>-</sup> cells from HD (grey), memory B cells from HD (green), memory B cells from ITP patients (blue) and residual memory B cells from RTX failure patients (red) according to the gating strategy shown in Figure 4. **(B)** Histograms showing MFIs (±SD) of CD32B, CXCR5, CD72, CD11c, FCRL4 and FCRL5 in HD (n=6), ITP (n=3) and RTX failure (n=7) patients. **(C)** Splenocytes from HD (n=3) and RTX Failure patients (n=3) were stimulated by CpG or CD40L and IL21/IL2/IL4/BAFF. Percentage of proliferating memory B cells was assessed after 5 days of culture with Cell proliferation Dye eFluor450. Two-tailed Mann-Whitney tests were performed ($***P < 0.001$; $**P < 0.01$, $*P < 0.05$).

| Patients | Age/Gender | ITP duration (months)<br>before splenectomy | Interval between RTX<br>and splenectomy<br>(months) | Response to RTX<br>(duration, months) | Treatments received before RTX<br>and splenectomy | Treatments received during<br>the month<br>preceding splenectomy | Response to<br>splenectomy |
| --- | --- | --- | --- | --- | --- | --- | --- |
| RTX_relapse1 | 30/F | 26 | 12 | CR (6 months) | CST, Ivlg | Ivlg | Failure |
| RTX_relapse2 | 61/F | 28 | 15 | CR (13 months) | CST, danazol | CST | CR |
| RTX_relapse3 | 42/F | 21 | 12 | CR (11 months) | CST | CST | CR |
| RTX_relapse4 | 51/M | 60 | 11 | CR (5 months) | CST, Ivlg | Ivlg | CR |
| RTX_relapse5 | 65/M | 15 | 11 | CR (6 months) | CST | CST | CR |
| RTX_relapse6 | 52/F | 44 | 8 | CR (6 months) | CST, Ivlg, danazol | CST, Ivlg | Failure |
| RTX_relapse7 | 45/F | 21 | 14 | CR (12 months) | CST, DAPS | CST | CR |
| RTX_relapse8 | 62/F | 156 | 14 | CR (12 months) | CST, TPO-RA, Ivlg, DAPS | TPO-RA, Ivlg | CR |
| RTX_failure1 | 59/F | 12 | 4 | Failure | CST, Ivlg | CST, Ivlg | CR |
| RTX_failure2 | 39/M | 360 | 4 | Failure | CST | CST | CR |
| RTX_failure3 | 22/F | 12 | 3 | Failure | TPO-RA, CST, Vcr | TPO-RA, CST | CR |
| RTX_failure4 | 24/M | 36 | 1 | Failure | CST, Vcr | CST, Vcr, tacrolimus | CR |
| RTX_failure5 | 48/F | 24 | 4 | Failure | CST, Ivlg, AZA | CST, Ivlg, AZA | Failure |
| RTX_failure6 | 51/F | 3 | 3 | Failure | CST, Ivlg, Vcr, danazol | CST, Ivlg, Vcr, danazol | CR |
| RTX_failure7 | 76/M | 23 | 9 | Failure | TPO-RA, CST, Ivlg | CST | Failure |
| RTX_failure8 | 54/M | 3 | 4 | Failure | CST, DAPS | CST, DAPS | Failure |
| RTX_failure9 | 66/F | 6 | 4 | Failure | CST, Ivlg, DAPS | CST, Ivlg | CR |
| RTX_failure10 | 27/M | 8 | 6 | Failure | CST, Ivlg, DAPS, HCQ | CST, Ivlg, DAPS, HCQ | CR |
| RTX_failure11 | 18/M | 6 | 2 | Failure | CST, Ivlg, TPO-RA, Vcr | CST, Ivlg, TPO-RA | CR |
| RTX_failure12 | 36/F | 6 | 3 | Failure | CST, Ivlg, TPO-RA, Vcr, AZA | CST, Ivlg, TPO-RA, Vcr | CR |
| RTX_failure13 | 28/F | 36 | 6 | Failure | CST, Ivlg, DAPS, Vcr | CST, Ivlg, DAPS, Vcr | CR |
| RTX_failure14 | 74/F | 48 | 6 | Failure | CST, Ivlg, DAPS | CST, Ivlg, DAPS | Failure |
| RTX_failure15 | 67/M | 6 | 6 | Failure | CST, Ivlg, TPO-RA, danazol, MMF | CST, Ivlg | CR |
| RTX_failure16 | 37/M | 60 | 1 | Failure | CST, Ivlg, TPO-RA, CSA, MMF | CST, Ivlg, TPO-RA, CSA, MMF | CR |
| ITP1 | 25/F | 84 | NA | NA | CST, Ivlg, DAPS | CST | CR |
| ITP2 | 40/F | 12 | NA | NA | IgIV, AZA, CST, DAPS | IgIV, AZA | CR |
| ITP3 | 60/H | 36 | NA | NA | CST, Ivlg | IgIV | CR |
| ITP4 | 50/F | 28 | NA | NA | CST, TPO-RA | TPO-RA | CR |
| ITP5 | 58/M | 144 | NA | NA | CST, Ivlg, TPO-RA | Ivlg, TPO-RA | CR |
| ITP6 | 22/M | 24 | NA | NA | CST | CST | Failure |
| ITP7 | 18/F | 12 | NA | NA | CST | CST | CR |
| HD1 | 52/F | NA | NA | NA | NA | NA | NA |
| HD2 | 22/M | NA | NA | NA | NA | NA | NA |
| HD3 | 32/F | NA | NA | NA | NA | NA | NA |
| HD4 | 28/M | NA | NA | NA | NA | NA | NA |
| HD5 | 62/M | NA | NA | NA | NA | NA | NA |
| HD6 | 64/M | NA | NA | NA | NA | NA | NA |
| HD7 | 90/F | NA | NA | NA | NA | NA | NA |
| HD8 | 68/M | NA | NA | NA | NA | NA | NA |
| HD9 | 60/F | NA | NA | NA | NA | NA | NA |

AZA : Azathioprine, CSA : Ciclosporine A, CST: Corticosteroids, Ivlg : Intraveinuous immunoglobulins, DAPS : Dapsone, HCQ : Hydroxychloroquine, MMF : Mycophenolate mofetil, TPO-RA : Thrombopoietine receptor agonists, RTX: Rituximab

Supplementary Table 1: Patients characteristics

| Patient | Population | Number of sorted cells | Isotype | initial reads | submitted IMGT | Number of productive sequences | Number of unique sequences | number of clones | d50 |
| --- | --- | --- | --- | --- | --- | --- | --- | --- | --- |
| RTX relapse 1 | Naive | 100000 | MU | 113836 | 7912 | 17975 | 7690 | 7579 | 2775 |
|  | Memory | 130000 | GAMMA | 104892 | 2773 | 19307 | 2535 | 1167 | 69 |
|  |  |  | MU | 105582 | 5833 | 24859 | 5509 | 3042 | 35 |
|  | GC | 70000 | GAMMA | 101734 | 3058 | 15481 | 2861 | 1294 | 82 |
|  |  |  | MU | 105943 | 4082 | 23315 | 3394 | 1336 | 50 |
|  | PC | 70000 | GAMMA | 172413 | 1847 | 5820 | 1817 | 1221 | 142 |
|  |  |  | MU | 110724 | 3119 | 16658 | 3072 | 1546 | 101 |
| RTX relapse 2 | Naive | 100000 | MU | 107764 | 7892 | 16734 | 7630 | 7518 | 2763 |
|  | Memory | 200000 | GAMMA | 119216 | 5991 | 20012 | 5581 | 3625 | 61 |
|  |  |  | MU | 103684 | 5814 | 24496 | 5216 | 2951 | 203 |
|  | GC | 10000 | GAMMA | 116882 | 2676 | 23672 | 2484 | 575 | 24 |
|  |  |  | MU | 102299 | 3327 | 34079 | 3123 | 1042 | 21 |
|  | PC | 50000 | GAMMA | 147534 | 2223 | 8985 | 2155 | 1286 | 60 |
|  |  |  | MU | 96436 | 2784 | 11445 | 2758 | 2166 | 391 |
| RTX relapse 3 | Naive | 100000 | MU | 1410223 | 50521 | 119460 | 45552 | 38568 | 9259 |
|  | Memory | 30000 | GAMMA | 1302391 | 17385 | 92280 | 15358 | 5157 | 33 |
|  |  |  | MU | 1481840 | 16294 | 80997 | 13508 | 5448 | 349 |
|  | GC | 20000 | GAMMA | 1384055 | 22692 | 88691 | 19497 | 5806 | 217 |
|  |  |  | MU | 1600195 | 30613 | 160998 | 26058 | 6523 | 104 |
|  | PC | 80000 | GAMMA | 1427490 | 12732 | 31536 | 11805 | 7002 | 451 |
|  |  |  | MU | 1766698 | 35347 | 163650 | 33758 | 10485 | 288 |

Supplementary Table 2: Immunoglobulin heavy chain sequencing

|  | GC | Memory | PC |
| --- | --- | --- | --- |
| RTX relapse 1 | 1.5295 | 1.9508 | 2.8172 |
| RTX relapse 2 | 1.5959 | 2.2332 | 2.0971 |
| RTX relapse 3 | 3.6396 | 2.2199 | 3.7895 |

**Supplementary Table 3: Mean standard deviation of VH mutation number in the first 100 clones**

| Patients | Subset | Number of sorted cells | Number of IgG secreting clones | Number of GPIIbIIIa specific clones | Percentage of GPIIbIIIa specific clones among IgG secreting clones |
| --- | --- | --- | --- | --- | --- |
| RTX_relapse1 | GC | 1152 | 214 | 47 | 21.96 |
| RTX_relapse1 | IgG+ Memory | 1152 | 438 | 51 | 11.64 |
| RTX_relapse2 | GC | 1440 | 89 | 31 | 34.83 |
| RTX_relapse2 | IgG+ Memory | 576 | 236 | 61 | 25.85 |
| RTX_relapse3 | GC | 1632 | 190 | 27 | 14.21 |
| RTX_relapse3 | IgG+ Memory | 1248 | 430 | 68 | 15.81 |
| RTX_relapse4 | GC | 1056 | 261 | 41 | 15.71 |
| RTX_relapse4 | IgG+ Memory | 672 | 307 | 42 | 13.68 |
| RTX_relapse5 | GC | 480 | 84 | 0 | 0.00 |
| RTX_relapse5 | IgG+ Memory | 480 | 113 | 0 | 0.00 |
| RTX_relapse6 | GC | 576 | 36 | 7 | 19.44 |
| RTX_relapse6 | IgG+ Memory | 576 | 241 | 38 | 15.77 |
| RTX_relapse7 | GC | 576 | 90 | 3 | 3.33 |
| RTX_relapse7 | IgG+ Memory | 576 | 127 | 1 | 0.79 |
| RTX_relapse8 | GC | 576 | 0* | 0 | ND |
| RTX_relapse8 | IgG+ Memory | 576 | 0* | 0 | ND |
| RTX_failure1 | Residual Memory | 1152 | 179 | 27 | 15.00 |
| RTX_failure2 | Residual Memory | 1152 | 43 | 2 | 4.70 |
| RTX_failure3 | Residual Memory | 768 | 124 | 13 | 11.00 |
| RTX_failure4 | Residual Memory | 1152 | 134 | 16 | 12.00 |
| HD5 | GC | 576 | 89 | 3 | 3.37 |
| HD5 | IgG+ Memory | 288 | 157 | 1 | 0.64 |
| HD6 | GC | 576 | 80 | 0 | 0.00 |
| HD6 | IgG+ Memory | 576 | 230 | 0 | 0.00 |
| HD8 | GC | 576 | 227 | 9 | 3.96 |
| HD8 | IgG+ Memory | 288 | 183 | 3 | 1.64 |

**Supplementary Table 4: Identification of anti-GPIIbIIIa specific single B cells**

| Antigen | Fluorochrome | Origin | Clone |
| --- | --- | --- | --- |
| CD38 | PerCP Cy5.5 | Sony | HIT2 |
| CD38 | Brilliant Blue 700 | BD Biosciences | HIT2 |
| CD27 | APC | Sony | MT271 |
| CD27 | PE-Cy7 | BD Biosciences | LG.3A10 |
| CD19 | PECF594 | BD | HIB19 |
| CD19 | Alexa Fluor 700 | BD Biosciences | HIB20 |
| CD24 | FITC | BD | ML5 |
| CD24 | PECy7 | Sony | ML5 |
| CD24 | FITC | BD Biosciences | ML5 |
| CD21 | PECy7 | BD | BLY4 |
| CD20 | APC H7 | BD | 2H7 |
| CD3 | af700 | BD | UCHT1 |
| CD3 | Brilliant Ultraviolet 395 | BD Biosciences | UCHT1 |
| CD3 | PE | BD Biosciences | UCHT1 |
| CD14 | af700 | BD | MφP9 |
| CD14 | Brilliant Ultraviolet 395 | BD Biosciences | MφP9 |
| CD14 | PE | BD Biosciences | M5E2 |
| CD16 | Brilliant Ultraviolet 395 | BD Biosciences | 3G8 |
| CD16 | PE | BD Biosciences | 3G8 |
| CD16 | af700 | BD | 3G8 |
| CD10 | PE | BD | Hi10a |
| IgD | bv421 | BD | IA6-2 |
| IgD | Brilliant Violet 711 | BD Biosciences | IA6-2 |
| IgD | PECF594 | BD Biosciences | IA6-2 |
| IgM | bv 605 | Biolegend | MHM88 |
| IgM | Brilliant Violet 510 | BD Biosciences | G20-127 |
| IgG | Brilliant violet 605 | BD Biosciences | G18-145 |
| IgG | APC H7 | BD | G18-145 |
| IgA | APC | Miltenyi | IS11-8E10 |
| IgA | PE | Miltenyi | IS11-8E10 |
| Ki67 | af488 | BD | B56 |
| FcγRIIb (CD32b) | Alexa 488 | Gift from Dr P. Bruhns | N297A |
| FcRL4 (CD307d) | PE | BD Biosciences | 413D12 |
| CD72 | FITC | BD Biosciences | J4-117 |
| FcRL5 (CD307e) | PE | BD Biosciences | 509F6 |
| CD22 | APC | BD Biosciences | HIB22 |
| CXCR5 | Brilliant Blue 515 | BD Biosciences | RF8B2 |
| CD79b | PE | BD Biosciences | CB3-1 |
| pY759 -PLCy2 | Alexa 488 | BD Biosciences | K86-689.37 |
| pY84-BLNK | PE | BD Biosciences | J117-1278 |
| Live Dead Aqua |  | ThermoFisher |  |
| Zombie violet |  | Biolegend |  |
| Sytox Blue |  | ThermoFisher |  |

**Supplementary Table 5: Antibodies and clones**

| NGS Primers |  |
| --- | --- |
| dsDNA Primers |  |
| L_VH1*-r | CTCGGAGATGTGTATAAGAGACAGNNNNNNNNNNNNNNNNCAACTACAGGTGCCCACTCC |
| L_VH1-46-r | CTCGGAGATGTGTATAAGAGACAGNNNNNNNNNNNNNNNNTAGCTCCAGGTGCTCACTCC |
| L_VH1-69-r | CTCGGAGATGTGTATAAGAGACAGNNNNNNNNNNNNNNNNCAGCYACAGGTGTCCASTCC |
| L_VH1-2-r | CTCGGAGATGTGTATAAGAGACAGNNNNNNNNNNNNNNNNCMACAGGWGCCCACTCC |
| L_VH1-45-r | CTCGGAGATGTGTATAAGAGACAGNNNNNNNNNNNNNNNNAGCCACAGATGCCTACTCC |
| L_VH1-24-r | CTCGGAGATGTGTATAAGAGACAGNNNNNNNNNNNNNNNNCTACAGGCACCCACGCC |
| L_VH2-r | CTCGGAGATGTGTATAAGAGACAGNNNNNNNNNNNNNNNNCCKTCCTGGGTCTTRTCC |
| L_VH2-70*09-r | CTCGGAGATGTGTATAAGAGACAGNNNNNNNNNNNNNNNNCCTTCATGGGTCTTGTCT |
| L_VH3*-r | CTCGGAGATGTGTATAAGAGACAGNNNNNNNNNNNNNNNNTTWAAAGGTGTCCAGTGTGARG |
| L_VH3-30/33/11-r | CTCGGAGATGTGTATAAGAGACAGNNNNNNNNNNNNNNNNWTAARAGGTGTCCAGTGTCAGG |
| L_VH4-r | CTCGGAGATGTGTATAAGAGACAGNNNNNNNNNNNNNNNNCCAGATGGGTCCTGYCC |
| L_VH5-51-r | CTCGGAGATGTGTATAAGAGACAGNNNNNNNNNNNNNNNNNTTCTCCAAGGAGTCTGTGCC |
| L_VH6-1-r | CTCGGAGATGTGTATAAGAGACAGNNNNNNNNNNNNNNNNCCATGGGGTGTCCTGTCA |
| PCR1 primers |  |
| Forward primer |  |
| IL-R2 | GTCTCGTGGGCTCGGAGATGTGTATAAGAGACAG |
| Reverse primers |  |
| CHM-2DW | CAGGAGACGAGGGGGAAAAGG |
| CHA-2DW | GGAAGAAGCCCTGGACCAGGC |
| CHG-D1 | TTCGGGGAAGTAGTCCTTG |
| CHD-n1 | TATCAAGCATGCCAGGACCAC |
| PCR2 primers |  |
| Reverse primers |  |
| HCA-n2p_ILr | TCGTCGGCAGCGTCAGATGTGTATAAGAGACAGGCGATGACCACGTTCCCATCT |
| HCGn2p_ILr | TCGTCGGCAGCGTCAGATGTGTATAAGAGACAGGAAGTAGTCCTTGACCAGGCA |
| HCM-n2m_ILr | TCGTCGGCAGCGTCAGATGTGTATAAGAGACAGAAAGGGTTGGGGCGGATGC |
| HCD-n2m_ILr | TCGTCGGCAGCGTCAGATGTGTATAAGAGACAGGGGGAACACATCCGGAGCCT |
| Single cell primers |  |
| Cy CH1 | GGAAGGTGTGCACGCCGCTGGTC |
| L-VH mix: |  |
| L-VH1 | ACAGGTGCCCACTCCCAGGTGCAG |
| L-VH 3 | AAGGTGTCCAGTGTGARGTGCAG |
| L-VH 4/6 | CCCAGATGGGTCCTGTCCCAGGTGCAG |
| L-VH 5 | CAAGGAGTCTGTTCCGAGGTGCAG |

**Supplementary Table 6: Primers**
